# Dysregulated Proline Metabolism Contributes to Subretinal Fibrosis in Neovascular AMD: Therapeutic Potential of Prolyl-4-Hydroxylase Inhibition

**DOI:** 10.1101/2025.04.13.648057

**Authors:** Yue Zeng, Ting Zhang, Elisa Cornish, So-Ra Lee, Michelle Yam, Mark Eminhizer, Jingwen Zeng, Jialing Zhang, Shaoxue Zeng, Xian Wei, Jinliang Yang, Meidong Zhu, Andrew Chang, Meixia Zhang, Jianhai Du, Ling Zhu, Mark C. Gillies

## Abstract

Subretinal fibrosis, a major cause of irreversible vision loss in neovascular age-related macular degeneration (nAMD), is driven by excessive deposition of extracellular matrix such as collagens. While proline metabolism is known to play a critical role in collagen biosynthesis and fibrosis, its involvement in subretinal fibrosis remains unclear. Here, we characterized the progression of fibrovascular lesions in JR5558 mice, observing significant molecular alterations as early as 4 weeks of age and phenotypic changes by 8 weeks. Transcriptomic and metabolomic analyses revealed elevated levels of 4-hydroxyproline, an essential component of collagen, alongside significant alterations of other fibrosis-related pathways. P4HA1, a catalytic subunit of prolyl-4-hydroxylase essential for 4-hydroxyproline biosynthesis, was prominently expressed in fibrotic lesions in retinas of JR5558 and two-stage laser-induced murine models, as well as human eyes with nAMD. Targeting P4HA1 with the small-molecule inhibitor diethyl pythiDC significantly attenuated fibrovascular lesion growth in the JR5558 murine models and reduced collagen turnover in human retinal pigment epithelium cells. Combining diethyl pythiDC with aflibercept had a stronger antifibrotic effect than monotherapies in JR5558 mice. These findings suggest a key contribution of proline metabolism, particularly proline hydroxylation, in subretinal fibrosis. Inhibiting P4HA1 with diethyl pythiDC inhibited fibrosis in the models we studied, offering a novel therapeutic strategy. Further research is warranted to explore the potential benefits of combining existing anti-angiogenic therapies with drugs that inhibit proline metabolism for the management of nAMD-associated fibrosis.

## Introduction

Age-related macular degeneration (AMD) is the leading cause of blindness in the elderly population[1, 2]. Neovascular AMD (nAMD), accounting for 90% of AMD-associated vision loss, is characterized by the invasion of choroidal or retinal neovascularization in the macular region[3]. In nAMD, enhanced expression of vascular endothelial growth factor (VEGF) promotes chronic inflammation and uncontrolled wound healing, which ultimately results in subretinal scars. Despite anti-VEGF treatment as the first-line therapy for nAMD, large clinical trials have shown that the incidences of subretinal fibrosis have been reported in 26-46% within 2 years, 38-56% within 5 years and 50-70% within 10 years[4, 5]. Early histological studies of nAMD-affected human eyes revealed that subretinal fibrotic lesion thickness correlated directly with photoreceptor degeneration[6]. Clinical studies have highlighted that subretinal fibrosis is a major cause of long-term poor visual outcomes in patients with nAMD[7, 8]. However, no US Food and Drug Administration-approved interventions currently exist to prevent or treat this irreversible complication.

Excessive deposition of extracellular matrix (ECM) proteins from Müller cells, retinal pigment epithelium (RPE) cells, endothelial cells, pericytes and macrophages is the hallmark of subretinal fibrosis[9–12]. Collagens, characterized by triple-helical structures, are the main structural proteins found in ECM[13]. Recent studies have highlighted the emerging role of collagen metabolic reprogramming in the pathogenesis of fibrosis, particularly the biosynthesis of its essential building blocks, such as glycine and proline[14–16]. Hydroxyproline, which stabilizes collagen’s triple helix at physiological temperature, is derived from the post-translational modification of proline residues. This process is catalyzed by prolyl-4-hydroxylase (P4H). Three forms of P4H have been reported to date, including hypoxia-inducible transcription factor P4H (HIF-P4H), collagen P4H (C-P4H) and transmembrane P4H (P4H-TM). C-P4H, hereafter referred to as P4H, is directly involved in collagen synthesis and thus, collagen-related disorders[17]. P4H is an α_2_β_2_ tetramer comprising of two identical α-subunits (encoded by *P4HA1*, *P4HA2*, or *P4HA3*) and two structural β-subunits (encoded by *P4HB*) [18]. This hydroxylation process in collagen is one of the most abundant metabolic fluxes in the human body[19]. Elevated P4H expression has been reported in tissues from patients with cholestatic liver injury[20] and idiopathic pulmonary fibrosis (IPF)[21], as well as in response to various pro-fibrotic factors such as TGFβ, FGF2 and EGF[22–24]. As the main α isoform in most cells, P4HA1 knockout caused embryonic lethality in mice due to the abnormal assembly of collagens[25]. Several studies have shown P4HA1 as a downstream target for fibrosis-related diseases[26–28]. However, the potential role of P4HA1 in ocular fibrosis has been unclear.

The hydroxylation reaction catalyzed by P4H involves two distinct stages. In the first stage, α-ketoglutarate (αKG) undergoes oxidative decarboxylation to produce succinate and a highly reactive Fe (IV)=O species (ferryl ion). In the second stage, this ferryl ion facilitates the hydroxylation of proline residues on collagen peptides through a radical rebound mechanism, completing the enzymatic reaction[29]. P4H-specific inhibitors have been widely developed and demonstrated to alleviate IPF, liver fibrosis and bladder outlet obstruction *in vivo*[21, 30, 31]. Diethyl pythiDC, an enzyme-specific αKG mimic, is the most potent inhibitor of P4H reported to date (Ki = 0.39 µM) and displays modest selectivity for human P4H[32]. Notably, it inhibited collagen biosynthesis in human breast cancer cells with negligible affinity to iron[32]. Though diethyl pythiDC has been recognized as a promising drug in the cancer field, its therapeutic role in fibrotic disease, particularly in subretinal fibrosis, remains poorly understood[33–36].

JR5558 transgenic mice have been characterized as a reliable model for studying both the early and late stages of nAMD, which are associated with neovascularization and subretinal fibrosis[37, 38]. Given that the distinct, yellow-white, subretinal fibrovascular lesions appear spontaneously in this model, their nature and progression may differ from those in other mouse models of subretinal fibrosis.

In the current study, we explored the transcriptional and metabolic profiles of this mouse model to elucidate the molecular cues governing this pathological process. We aimed to identify new insights into the role of aberrant metabolism in subretinal fibrosis, opening new avenues for the treatment of this irreversible disease. By comparing the transcriptomic and metabolic profiles of retinas from JR5558 and wild-type mice, coupled with *in vitro* and *in vivo* validations, we sought to explore the therapeutic role of blocking P4HA1 in subretinal fibrosis through inhibiting collagen metabolism.

## Materials and Methods

### Animals

All procedures in JR5558 mice and wild-type controls were approved by the University of Sydney Animal Ethics Committee (Approval No. 2021/2013). The procedures in mice receiving two-stage laser photocoagulation were approved by the Experimental Animal Ethics Committee of Sichuan University (Approval No. 20240925001). All mice experiments were performed following the Association for Research in Vision and Ophthalmology (ARVO) Statement for the Use of Animals in Ophthalmic and Vision Research. JR5558 mice with C57BL/6J background were purchased from the Jackson Laboratory (strain #:005558; RRID: IMSR_JAX:005558). The genotype was confirmed by PCR amplification with tail DNA samples. Wild-type C57BL/6J mice were used as controls or underwent two-stage laser photocoagulation. All animals were housed in the pathogen-free facilities in a 12 h light/dark cycle with free access to food and water.

### Two-stage laser photocoagulation

A two-stage laser photocoagulation was performed to induce the subretinal fibrosis in mice as previously described[11]. In brief, 8-week-old C57BL/6J mice were anesthetized by intraperitoneal injection of pentobarbital sodium (2% w/v) after dilating pupils with Tropicamide Phenylephrine Eye Drops (Mydrin-P). Three laser burns were applied in each eye at 3, 6 and 9 o’clock positions of the fundus, approximately one disc diameter from the optic nerve using a 532nm Green Laser Photocoagulator (MERILAS 532a). Laser settings were: 250 mW power, 0.1 s duration and 100 μm spot size. Seven days later, a second laser burn was placed at the same area with the same laser settings.

### In vivo imaging

Color fundus images, optical coherence tomography (OCT) and fundus fluorescein angiography (FFA) were captured using the Phoenix MICRON IV Retinal Imaging System (Phoenix Technology Group, USA). Mice were anaesthetized, followed by pupil dilation and cornea lubrication as described above. For fundus images, the optic nerve was centered and the location of retinal vasculatures was aligned during each follow-up to ensure a consistent area for quantification. Ten cross-sectional OCT images were captured per eye at 15 μm intervals above and below the optic nerve head. For FFA, animals were subcutaneously injected with 100mg/mL fluorescein sodium (SERB, France) and images of the early and mid-to-late phases were captured at 1 minute and 5 minutes after injection, primarily focusing on the left eye. After imaging, mice were recovered from anesthesia by atipamezole (1.2mg/kg, Troy Laboratories, Australia) on a heating mat and returned to the cage.

### Quantification of fundus images

Lesion areas in the fundus photograph of JR5558 mice were quantified using ImageJ software (http://imagej.nih.gov/ij/, National Institutes of Health, USA) and a custom Macro (**Supplementary file S1**) was developed to ensure consistent and reproducible analysis during each follow-up. In detail, a circular region of interest (ROI) with a radius of 283 μm centered on the optic nerve head was created, covering approximately 50% of the total fundus area. The image was then converted to an 8-bit grayscale format and the background was removed using a 200 μm rolling ball algorithm. Lesions fully or partially located within the ROI were selected using the freehand selection tool and refined by threshold adjustment. Lesion areas were measured using the particle analysis in ImageJ and results were cross-checked with the original fundus images. The total lesion area for each eye was calculated for each follow-up. Each image was analyzed by two investigators masked to the treatment group and the average value was used for further analysis.

FFA was used as a reference to classify lesions in fundus photographs. Lesion leakage in FFA was graded as follows: Grade 1, no significant leakage or mottled hyperfluorescence; Grade 2, hyperfluorescent lesion without a progressive increase in size or intensity; Grade 3, hyperfluorescent lesion with an increase in size and/or intensity [39, 40]. Lesions with grade 1 or 2 leakage were defined as primarily fibrotic lesions and the total area was subsequently quantified.

### Cell Culture

Human primary RPE cells (huRPEs) were cultured from RPE/choroid complexes dissected from donor eyes and digested in 7ml TrypLE^TM^ (12604013, Thermo Fisher Scientific) in a cell-culture incubator for 1 h. The tissues were removed from the solution and digestion was terminated by DMEM medium (10569, Gibco) supplemented with 10% fetal bovine serum (FBS, F9423, Sigma-Aldrich) and 1% penicillin-streptomycin (P/S, 4333, Sigma-Aldrich). Following centrifugation, cell pellets were resuspended in the complete medium (DMEM with 10% FBS and 1% P/S) and seeded into a T25 culture flask (156367, Nunc™). The culture medium was changed after 24 h to avoid contamination and routine culture was continued with medium changes twice a week. Only huRPEs at passage 2 or earlier were included in subsequent experiments. All cells were maintained in a 37°C incubator with 5% CO2 and 95% humidity. HuRPEs were starved overnight with DMEM supplemented with 1% FBS and 1% P/S and treated with 12.5ng/mL TGFβ2 (302-B2/CF, R&D Systems) for 3 days to induce fibrotic features *in vitro*.

### AlamarBlue Cell Viability Assay

HuPMCs and huRPEs were seeded in 96-well plates (3300, Corning, NY). At 70% confluency, cells were starved overnight with DMEM supplemented with 1% FBS and 1% P/S. The following day, cells were treated with starvation medium containing different concentrations of diethyl pythiDC (3.125 μM, 6.25 μM, 12.5 μM, 25 μM, 50 μM, 100 μM; HY-103068, MedChemExpress) or 0.2% DMSO (D-2650, Sigma-Aldrich). After a 24-hour treatment, the medium was replaced with starvation medium containing 10% AlamarBlue reagent (DAL1100, Thermo Fisher Scientific), followed by a 4-hour incubation at 37°C. The fluorescence was detected by a plate reader (FLUOstar Omega, BMG Labtech, Germany) according to the manufacturer’s protocol and values were normalized to the DMSO control group.

### *In vivo* treatment

Diethyl pythiDC was dissolved in DMSO at a concentration of 50mM. Following *in vivo* imaging, 4-week-old JR5558 mice were anaesthetized and intravitreally injected with 1 μL of 100 μM diethyl pythiDC or vehicle control, 0.8% DMSO in both eyes. For the combination therapy, four-week-old JR5558 mice were intravitreally injected with 1 μL of 100 μM diethyl pythiDC, 2.5µg/µL aflibercept, 2.5µg/µL aflibercept combined with 100µM diethyl pythiDC, or 0.8% DMSO in both eyes. Fundus photograph and OCT were captured 2 weeks, 4 weeks and 8 weeks after the injection. FFA was only performed 8 weeks after the injection.

### Immunofluorescence of frozen sections and mouse RPE/choroid flatmounts

Human retinas were obtained from post-mortem healthy donors, with consent and ethical approval from Human Research Ethics Committee of the University of Sydney (Protocol Numbers: 2016/282) and Sydney Local Health District Ethics Review Committee (2020/ETH03339). For frozen sections, post-mortem eyes from an nAMD patient and enucleated eyes from 8-week-old JR5558 mice, age-matched control mice and mice at day 30 after the second laser were fixed in 4% paraformaldehyde (PFA, C005, ProSciTech) for 1h at room temperature (RT), followed by washing with fresh PBS (09-2051-100, Medicago). Eyes were equilibrated at 30% sucrose overnight and embedded in O.C.T. Compound (IA012, Scigen). The frozen blocks were sectioned at 10 μm thickness for mice eyes and 16 μm for nAMD eyes using a CryoStat (LEICA CM3050S, Germany). Cryosections were then blocked with PBS containing 5% donkey serum (566460, Millipore) and 1% Triton X-100 (108603, Merck) for 1 h at RT. After blocking, cryosections were incubated with primary antibodies at 4°C for 48-72 hours. Alexa Fluor 488- and 594-conjugated secondary antibodies were incubated the next day for 3 h at RT. The primary and secondary antibodies were diluted in 1% donkey serum and Triton X-100 (0.5% for cells, 1% for cryosections) in PBS. Samples were counterstained with Hoechst 33258 (H3569, Thermo Fisher Scientific) for 5 min at RT, mounted in Vectashield Antifade Mounting Medium (H1000, Vector Laboratories, USA) and cover-slipped.

For mouse RPE/choroid flatmounts, the orientation of the JR5558 mouse eye was ensured by marking the superior area of the sclera before performing retinal imaging. Eyes enucleated at 8 weeks of age or 8 weeks after the intravitreal injection were fixed in 4% PFA for 1h at RT. Eyes were then immersed in PBS before anterior segments and neural retina were carefully removed. During dissection, 1 mm incision was made along the marker to visualize the upright orientation. RPE/choroid complexes were then blocked in 96-well plate (3598, Corning, NY) with 5% donkey serum and 1% Triton X-100 for 1h at RT, followed by incubation with primary antibody against Collagen I and Alexa Fluor 594-conjugated Isolectin-B4 (IB4) at 4°C for 48h. Samples were incubated with an Alexa Fluor 488-conjugated secondary antibody for 2h at RT, counter-stained with Hoechst 33258 for 15 min at RT, mounted and cover-slipped. The primary and secondary antibodies were diluted as in mouse cryosection staining. Fluorescence in all samples was imaged using a Zeiss LSM 900 Confocal Microscope (Zeiss, Germany). The antibodies used are listed in **Table 1**.

**Table 1:** Antibodies used for western blot (WB) and immunofluorescent (IF) studies.

| Antibody | Company | Source | Catalog No. | Working dilution |  |
| --- | --- | --- | --- | --- | --- |
|  |  |  |  | WB | IF |
| P4HA1 | Novus Biologicals | Goat | NB100-57852 | N/A | 1: 100 |
| P4HA1 | Proteintech | Rabbit | 12658-1-AP | 1: 1000 | N/A |
| P4HA1 | Novus Biologicals | Rabbit | NBP1-84398 | 1: 1000 | N/A |
| CRALBP | Abcam | Mouse | ab15051 | N/A | 1: 500 |
| CRALBP | Abcam | Rabbit | ab243664 | N/A | 1: 500 |
| GFAP | Dako | Rabbit | Z0334 | N/A | 1: 500 |
| RPE65 | Novus Biologicals | Mouse | NB-100-355 | N/A | 1: 200 |
| Collagen I | Abcam | Rabbit | ab34710 | N/A | 1:100 |
| Collagen I | Abcam | Rabbit | ab255809 | 1: 1000 | N/A |
| Collagen I | Abcam | Rabbit | ab138492 | 1: 1000 | N/A |
| IB4-594 | Invitrogen | - | I21413 | N/A | 1:50 |
| Fibronectin | Chemicon | Rabbit | ab2033 | 1:1000 | 1:100 |
| α-SMA | Cell Signaling Technology | Mouse | 48938 | 1: 1000 | 1: 200 |
| CTGF | Abcam | Rabbit | ab125943 | 1: 1000 | N/A |
| LOXL3 | LS Bio | Rabbit | LS-C165846 | 1: 1000 | N/A |
| GAPDH | Cell Signaling Technology | Rabbit | 2118 | 1: 2000 | N/A |
| β-Actin | Cell Signaling Technology | Rabbit | 4967 | 1: 2000 | N/A |
| $\alpha/\beta$-Tubulin | Cell Signaling Technology | Rabbit | 2148S | 1: 2000 | N/A |
| Anti-rabbit IgG | Cell Signaling Technology | Goat | 7074 | 1: 5000 | N/A |
| Anti-mouse IgG | Cell Signaling Technology | Horse | 7076 | 1: 5000 | N/A |
| Anti-mouse-488 | Invitrogen | Donkey | A-21202 | N/A | 1: 1000 |
| Anti-rabbit-488 | Invitrogen | Donkey | A-21206 | N/A | 1: 1000 |
| Anti-mouse-594 | Invitrogen | Donkey | A-21203 | N/A | 1: 1000 |
| Anti-rabbit-594 | Invitrogen | Donkey | A-21207 | N/A | 1: 1000 |
| Anti-goat-488 | Invitrogen | Donkey | A-11055 | N/A | 1: 1000 |

### Vibratome staining

Neural retinas from post-mortem human eyes were isolated as previously described[41]. A 10 × 20 mm rectangular piece of retinal tissue, spanning 4 mm nasally to 16 mm temporally from the fovea, was cut and placed in a 35 mm culture dish (430165, Corning) containing PBS. 4% PFA was used to fix the *ex vivo* retinal tissues for 1h at RT. After washing with PBS three times, samples were embedded in 3% low-melting agarose (50100, Lonza, Australia) in PBS. The blocks were then sectioned at 100 μm thickness using a vibratome (VT1200S, Leica, Germany) and placed in 48-well plates (3548, Corning, NY) with PBS at 4°C. Vibratome sections were blocked with 5% donkey serum and 1% Triton X-100 in PBS at 4°C overnight, then immunostained with primary antibodies against P4HA1 and CRALBP for 5 days at 4°C. Species-specific secondary antibodies conjugated with Alexa Fluor 488 and 594 were added for 2 days at 4°C, then counterstained with Hoechst for 30 min at RT. The dilution buffer for primary and secondary antibodies was the same as in cryosections staining. After mounting onto poly-lysine glass slides (22-042-941, Thermo Fisher Scientific), sections were sealed with adhesive spacers (S24737, Thermo Fisher Scientific) and captured with the Zeiss LSM 900 Confocal Microscope. The antibodies used are listed in **Table 1**.

### Protein Extraction and Western Blot

Cells were lysed in RIPA buffer containing 1% protease/phosphatase inhibitor cocktail (5872S, Cell Signaling). Tissue samples were homogenized on ice for 30 s in the same buffer and centrifuged at 12,000g, 4°C for 10 min. The supernatant was stored at −80°C until analysis. 400 μL medium from cells cultured in 24-well plates were collected and precipitated with 176 mg/mL ammonium sulfate (204501, Sigma-Aldrich). Tubes were vortexed and kept on a shaker at 4°C for 24 h. The mixture was centrifuged at 10,000g for 1 h at 4°C. The supernatant was discarded and pellets were dissolved in 40 μL RIPA buffer containing a 1% protease/phosphatase inhibitor cocktail for analysis. Protein concentration was measured using a BCA assay kit (QPBCA, Sigma-Aldrich). Total proteins mixed with NuPAGE loading dye (NP0007, Invitrogen) and reducing buffer (3483-12-3, Sigma-Aldrich) were heated at 70°C for 10 min. The same amount of proteins was loaded onto the NuPAGE 4-12% Tris-Bis gels (NP0001, Invitrogen) and electrophoresed at 180V for 70 min on ice. Gels were transferred onto polyvinylidene difluoride membrane (PVDF, Millipore) using the wet transfer system (#1703930, BioRad, USA). The PVDF membranes were blocked with 5% bovine serum albumin (BSA, A9417, Sigma-Aldrich) in Tris-buffered Saline (09-7500-100, Medicago) with 0.1% Tween 20 (BIO0777, Biochemicals) (TBST) for 1 h at RT, incubated with primary antibody overnight at 4°C and secondary antibody for 2 h at RT. Antibodies were diluted in 1% BSA in TBST and are listed in **Table 1**. The membranes were washed with TBST and then TBS for three times, incubated in ECL substrate (170-5061, Bio-Rad) for 5 min and imaged using a G-box imaging system (In Vitro Technologies). Bands were analyzed using the GeneTools image scanning and analysis package (Syngene, Cambridge). Results were quantified relative to loading controls (Tubulin or GAPDH) and presented as fold change relative to vehicle control groups.

### Collagen gel contraction assay

HuRPEs at 80% confluency were detached and suspended in DMEM complete medium. For each well, 50 μL type I collagen (3mg/mL, A1048301, Thermo Fisher Scientific), 12.5 μL 5X PBS, 37.5 μL cell suspension and 2 μL 1N NaOH were mixed thoroughly on ice in a 1.7 mL Eppendorf tube. The mixture was then transferred into a 96-well plate. Gels were polymerized in a cell culture incubator for 30 min before adding 100 μL DMEM starvation medium to each well. The following day, gels were dislodged from the edge of the well using a 25G needle. Medium was then replaced with DMEM starvation medium containing 12.5 ng/mL TGFβ2, 12.5 ng/mL TGFβ2 + 10 μM Diethyl pythiDC or vehicle controls and incubated for 3 days. Gel images were captured from day 1 to day 3 using a camera coupled to a Stereo Microscope (T2-HK830, AOSVI, China). Images were analyzed using ImageJ software (NIH, USA) to measure the gel area. Results were normalized to the gel area of vehicle control group or to day 0.

### RNA sequencing

Total RNA from the retinas of 4-week and 8-week-old JR5558 mice and age-matched C57BL/6 mice (n = 3) was extracted using a GenElute™ Single Cell RNA Purification Kit (RNB30, Sigma-Aldrich). RNA was quantified using a Qubit RNA Assay Kit (Q32852, Invitrogen) according to the manual. The quality control, library preparation and sequencing were commercially contracted to Novogene (https://www.novogene.com/us-en/). In brief, mRNA was purified through poly(A) selection using oligo(dT)-magnetic beads. After fragmentation, the first-strand cDNA was synthesized by using mRNA template and random hexamers primers, followed by the second-strand cDNA synthesis. The library was ready after end repair, A-tailing, adapter ligation, size selection, PCR amplification and purification. The qualified libraries were sequenced using Illumina NovaSeq 6000 (PE150) with paired end 150 bp read length.

### RNA data analysis

Raw reads were filtered to remove reads containing adapter, ploy-N and low-quality bases. The filtered clean reads were mapped to genome (GRCm39/mm39) using the HISAT2 v2.0.5 software. FeatureCounts v1.5.0-p3 was used to count the reads numbers mapped to each gene. The FPKM value for each gene was subsequently calculated using its length and the corresponding read count. Differential gene analysis using DESeq2 R package (1.20.0) was performed for pair-wise comparison between two groups. Genes with a p value < 0.05 was considered differentially expressed genes (DEGs). QIAGEN Ingenuity Pathway Analysis (IPA) was performed in genes with |log2FC| ≥ 1.3 and p value < 0.05. The license for IPA software was provided by the University of Sydney. Gene Ontology (GO) enrichment analysis of genes with |log2FC| ≥ 0.5 and p value < 0.05 and KEGG enrichment analysis of overlapped genes with p value < 0.05 were performed using the free online platform SRplot (https://www.bioinformatics.com.cn/en)[42].

### Metabolite analysis

Retinas from 4-week- and 8-week-old JR5558 mice and age-matched C57BL/6 mice were snap-frozen in dry ice and homogenized in 400 μL cold extraction buffer (methanol: chloroform: ddH2O, 700: 200: 50). 5 μL 0.1mg/mL Myristic-d27 acid were added to each sample as an internal standard. The tissue/extraction buffer mixture was incubated on ice for 20 minutes on a shaker, then centrifuged at 13,000g for 10 minutes at 4°C. The supernatant was collected in a clean tube and dried with a freeze dryer. The samples were then analyzed by gas chromatography–mass spectrometry (GC-MS) as previously reported[43, 44]. The ion abundance of each metabolite was normalized to the C57BL/6 controls to calculate the fold change. Partial least squares-discriminant analysis (PLS-DA) and variable importance in projection (VIP) were analyzed using MetaboAnalyst 6.0 (https://www.metaboanalyst.ca/)[45]. Pairwise ratios of metabolites within each donor were log2 transformed and calculated for fold change and p value in R (version 3.6.0).

### Statistical analysis

All data were presented as mean ± SEM. Statistical analysis was performed using GraphPad Prism 9.3.0 (GraphPad Software, San Diego, CA). Student’s t-tests were used to calculate the differences between the two groups. One-way ANOVA followed by Tukey’s or Dunnett’s post hoc test was applied for comparisons among three or more groups. K-means clustering of baseline lesion area was performed in Python (code available upon request). A p-value of < 0.05 was considered statistically significant.

## Results

### Characterization of fibrovascular changes in retinas from JR5558 mice

We first validated JR5558 mice as a mouse model of subretinal fibrosis by performing immunostaining with the vascular marker Isolectin B4 (IB4) and the fibrotic marker Collagen I in retinal cryosections (**Figure 1A**). Both markers were co-expressed in the subretinal lesion areas of the JR5558 mice retinas, confirming that the lesions were fibrovascular. Notably, we observed immunofluorescent staining for both CRALBP and RPE65, as well as co-staining of CRALBP and fibronectin in the subretinal lesions of JR5558 mice (**Figure S1**). These findings suggested that the fibrovascular lesions had components that originated from Müller cells and RPE cells. Western blot analysis revealed significant upregulated expression of fibrotic marker fibronectin and gliosis marker GFAP in 8-week-old JR5558 retinas, but not in 4-week-old JR5558 retinas (**Figure 1B-E**). To further characterize the fibrotic components within these lesions, RPE/choroidal flat mounts were stained with IB4 and Collagen I. Individual lesions were matched with corresponding fundus photography and FFA. We tried to define the different lesions through FFA together with the fundus photo by grading the leakage into three categories: grade 1, no significant leakage or mottled hyperfluorescence; grade 2, hyperfluorescent lesion without a progressive increase in size or intensity; and grade 3, hyperfluorescent lesion with an increase in size and/or intensity, as described in the Method section. We observed that lesions identified through FFA with grade 3 leakage (as defined in the Method) had a similar area of Collagen I and IB4 staining, while areas with grade 1 and 2 leakage had larger areas of Collagen I staining than IB4 staining (**Figure 1F**). Therefore, we used FFA as the reference to classify lesions with grade 1 and 2 leakage as primarily “fibrotic lesions” and calculated their sizes in fundus photos (**Figure 1G**). The size of fibrotic lesions increased significantly from 8 weeks onwards and stabilized at 12 weeks compared to 4 weeks of age (**Figure 1H**). Similar to a recent clinical observation, fibrotic scars in JR5558 mice appeared as subretinal hyperreflective material (SHRM) lesions on OCT with distinct boundaries, categorized into three types: Type A located underneath an intact RPE, type B located above the RPE, type C where the RPE was not distinguishable (**Figure 1I**). Quantitative analysis showed a significant increase in all lesion types from 8 weeks onwards (**Figure 1J**). Type A lesions were the most prevalent among all ages (**Figure 1K**).

**Figure 1:**
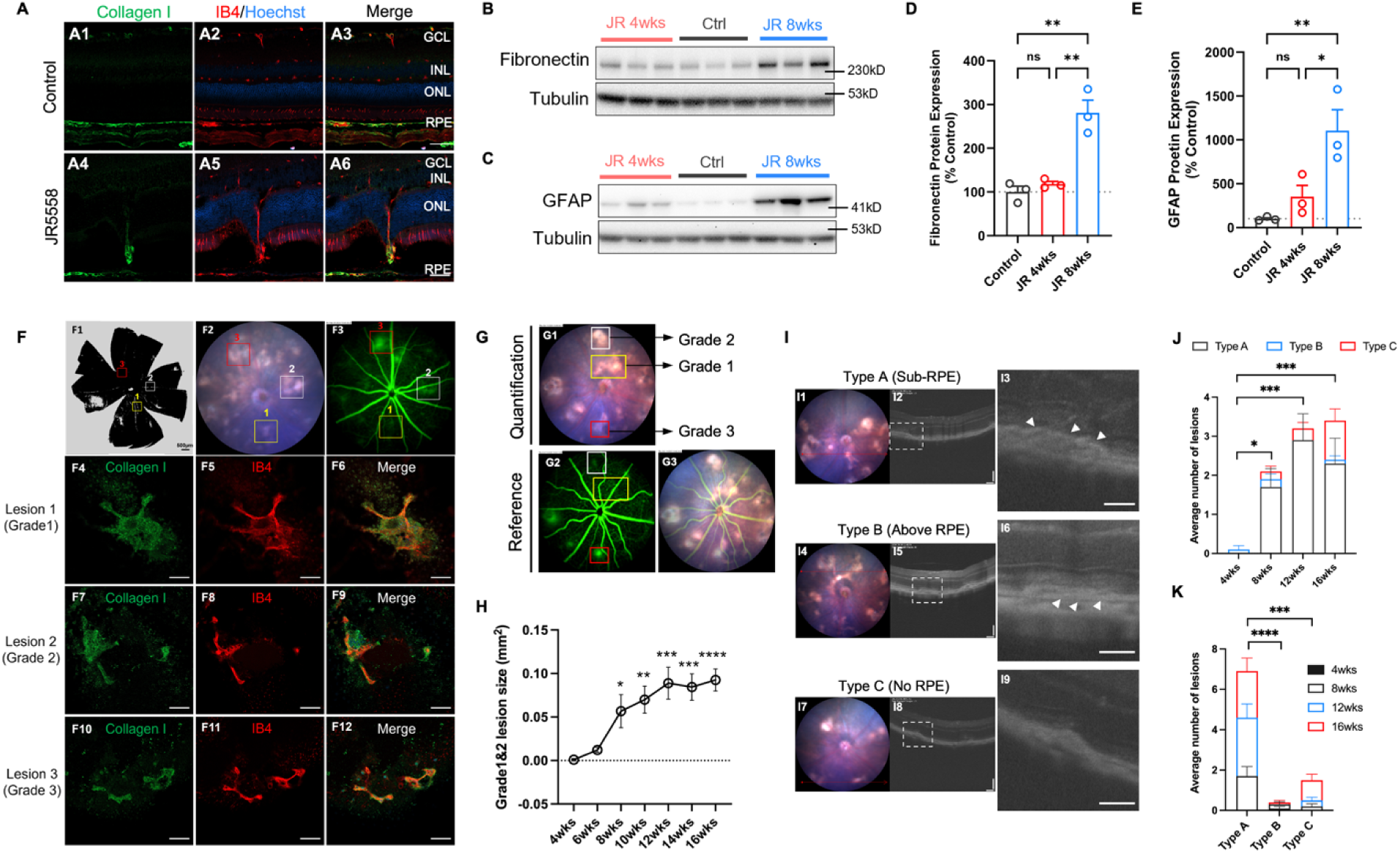
Phenotypic changes in JR5558 retinas appeared at 8 weeks of age. **(A)** Representative immunostaining of Collagen I (green) and Isolectin B4 (IB4) (red) in the retinas of 8-week-old control **(A1-A3)** and JR5558 **(A4-A6)** mice. Nuclei were stained with Hoechst (blue). Scale bar: 50 µm. **(B-E)** Western blots and quantification results showing protein expression of fibronectin **(B, D)** and GFAP **(C, E)** in retinas from 4- and 8-week-old JR5558 mice. Retinas from 8-week-old wild-type mice were used as controls. n = 3. **(F)** Representative staining of Collagen I (green) and IB4 (red) from JR5558 RPE/choroidal flatmounts **(F4-12**, scale bar: 50µm) with corresponding brightfield image **(F1,** scale bar: 500 µm), fundus photo **(F2)** and FFA **(F3)**. Yellow box: lesion 1, white box: lesion 2, red box: lesion 3. **(G)** Representative images showing methods for defining primarily “fibrotic lesion” (lesions with grade 1&2 leakage). **(H)** Statistical analysis of primarily “fibrotic lesion” areas quantified from fundus photos. *Indicates a statistically significant difference compared to lesion areas at 4 weeks of age. n = 11. **(I)** Fundus photo and corresponding OCT of three types of SHRM lesions. Magnified images of the lesion areas were shown in **(I3, I6, I9)**. Scale bar: 50 µm. **(J-K)** The average number of each lesion type at different ages in JR5558 mice. Data were shown as the average number of lesions per eye. n = 10. wks: weeks of age. GCL: ganglion cell layer. INL: inner nuclear layer. ONL: outer nuclear layer. RPE: retinal pigment epithelium. Statistical analysis was performed using one-way ANOVA followed by Tukey’s multiple comparison test in **(D-E)**, one-way ANOVA followed by Dunnett’s multiple comparison test in **(H)** and two-way repeated measures ANOVA followed by Tukey’s multiple comparison test in **(J-K)**. *: p < 0.05, **: p < 0.01, ***: p < 0.001, ****: p < 0.0001, ns: not significant. All data are presented as means ± SEM.

To check the molecular changes that may contribute to the development of fibrotic lesions, RNA sequencing was conducted on retinas from 4- and 8-week-old JR5558 and age-matched wild-type controls. Three retinas from three mice were collected and analyzed in each group (Figure 2A). Volcano plots showed 791 DEGs (405 upregulated, 386 downregulated) at 4 weeks and 841 DEGs (489 upregulated, 352 downregulated) at 8 weeks were significantly altered with |log2FC| ≥ 1.3 compared to controls (Figure 2B-C). Data reliability was confirmed by significant downregulation of *Crb1* gene at both ages[46]. We performed IPA analysis on these DEGs to identify the functional changes. The top 10 commonly changed pathways at both ages focused on fibrosis, complement system and acute phase response (Figure 2D-E). GO enrichment analysis highlighted collagen-containing extracellular matrix was the top enriched pathway at both ages (Figure 2F-G). Other fibrosis-related pathways, such as integrin binding[47, 48], wound healing[49] and cell adhesion molecule binding[50], were also significantly enriched in JR5558 retinas. Both IPA and GO analysis revealed significantly altered pathways in photoreceptors of JR5558 retinas, termed as phototransduction pathways, the visual cycle, retinoid binding and retinol binding. Damage to both rods and cones was confirmed by immunostaining in JR5558 retinal cryosections (**Figure S2**). Notably, JR5558 retinas showed clear upregulation of fibrosis-related genes starting from 4 weeks of age, including collagens, matrix-related genes, proteases, glycans and signaling molecules (Figure 2H-I). A Venn plot identified 1,291 significant DEGs that were consistently altered in JR5558 retinas at both ages compared to control retinas (Figure 2J). Further KEGG pathway analysis of these overlapped DEGs revealed that metabolic pathways ranked among the top 20 significantly enriched pathways, with arginine and proline metabolism showing the highest gene ratio (Figure 2K). The genes involved in this pathway were *p4ha1, amd2, ckmt1, gatm, lap3, aldh7a1* and *sat1*. Surprisingly, pathways related to angiogenesis, such as HIF-1 signaling pathway and VEGF signaling pathway, were enriched in the overlapped genes. Taken together, molecular cues indicating fibrosis were evident in JR5558 retinas as early as 4 weeks.

**Figure 2:**
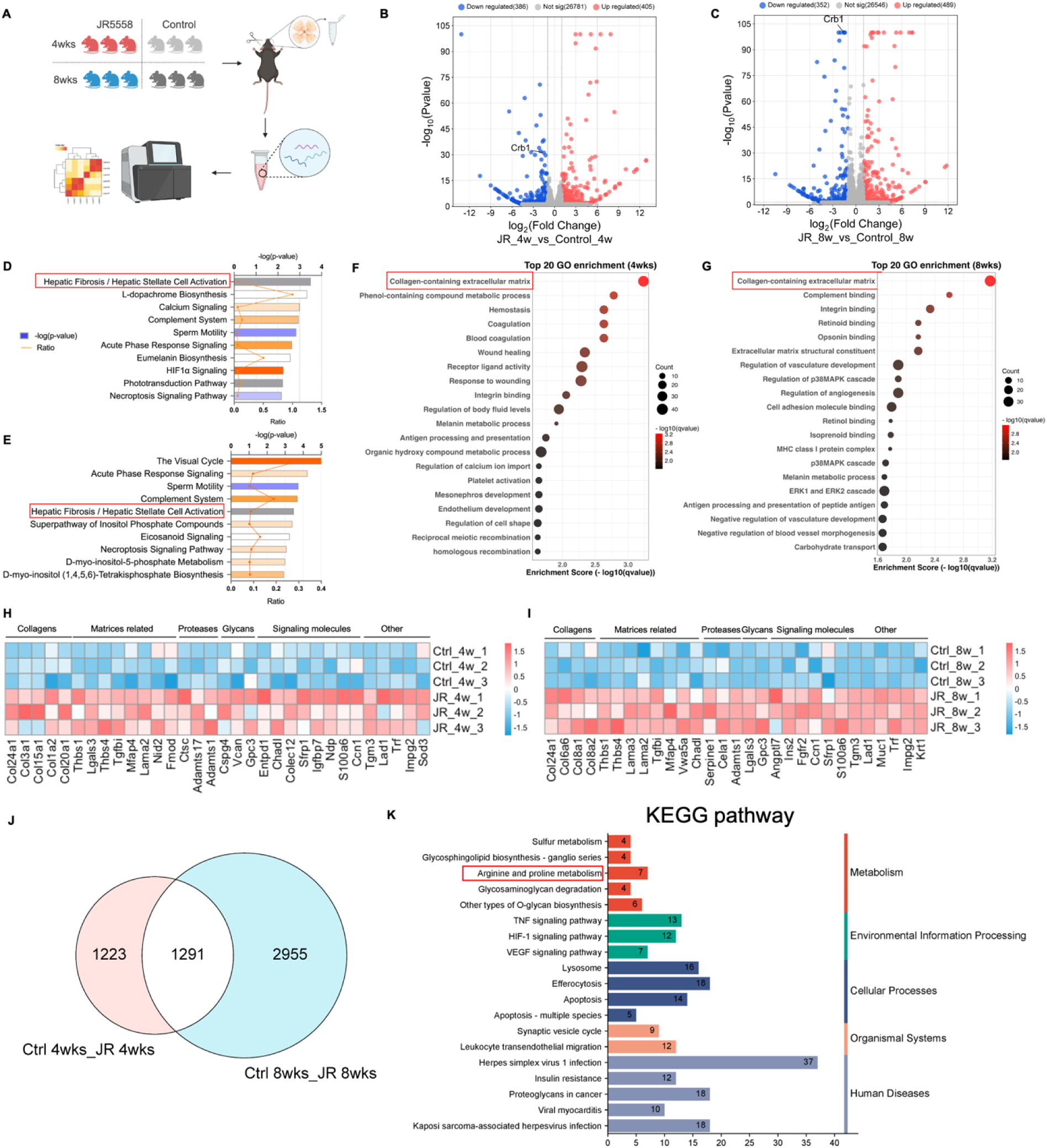
Transcriptomic changes of JR5558 mice occurred as early as 4 weeks of age. **(A)** Schematic diagram of experimental design of RNA-sequencing in JR5558 and control retinas. **(B-C)** Volcano plots showing DEGs between 4-week-old JR5558 vs control retinas **(B)**, 8-week-old JR5558 vs control retinas **(C)**. n = 3. **(D-E)** Ingenuity Pathway Analysis (IPA) analysis of DEGs between 4-week-old JR5558 versus control retinas **(D)**, 8-week-old JR5558 versus control retinas **(E)**. **(F-G)** Gene Ontology (GO) enrichment analysis of DEGs between 4-week-old JR5558 vs control retinas **(F)**, 8-week-old JR5558 vs control retinas **(G)**. **(H-I)**: Significantly altered fibrosis-related genes in 4-week-old JR5558 **(H)** and 8-week-old JR5558 **(I)** retinas compared to age-matched controls. **(J)** Venn plot comparing significant DEGs from “JR5558 4wks vs control 4wks” and “JR5558 8wks vs control 8wks”. **(K)** KEGG enrichment analysis of the 1,291 overlapped DEGs identified from **(J)**.

### Hydroxyproline was increased in JR5558 retinas

Given that both collagen-related components (Figure 2F**–G**) and collagen metabolic pathways (Figure 2J**– K**) were significantly enriched in JR5558 retinas based on transcriptomic analysis, we next examined whether collagen metabolism was functionally altered in this model by focusing on hydroxyproline, a key metabolite essential for collagen synthesis and stability. Four retinas per time point were collected from each 4- and 8-week-old JR5558 and control mice (Figure 3A). A total of 53 metabolites in amino acid and glucose metabolism were measured by GC-MS (**Supplementary file S2**). Partial Least Squares Discriminant Analysis (PLS-DA) illustrated clear separations in retinal metabolites between two groups at both ages, indicating the retinal metabolome of JR5558 retinas was significantly altered (Figure 3B-C). Variable importance in projection (VIP) scores were used to identify metabolites that are important for the separation in PLS-DA model. At both ages, 4-hydroxyproline was the only metabolite consistently ranked among the top five highest VIP scores, with a score of 1.653 at 4 weeks and 1.671 at 8 weeks (Figure 3D-E). 4-hydroxyproline has long been identified as a signature amino acid of fibrillar collagens, responsible for collagen thermal stability[18]. It was the top metabolite significantly increased in 4-week-old JR5558 retinas (2.82-fold, p=0.01). It showed higher relative increases at 8 weeks, with marginal statistical significance (4.67-fold, p=0.06) largely due to the inter-individual variation in one sample (Figure 3F-G). To reduce the impact of inter-group and experimental variability, we performed pairwise ratio comparisons for the 53 metabolites (**Supplementary file S3**). At both ages, 4-hydroxyproline ranked among the top 30 ratios with the most significant changes in JR5558 mice compared to controls (Figure 3H-I). Notably, 4-hydroxyproline was the most frequently observed numerator in pairwise ratio comparisons at 8 weeks (Figure 3J). These data indicated that dysregulated collagen metabolism, highlighted by elevated 4-hydroxyproline levels, was associated with subretinal fibrosis in JR5558 mice. We also identified significant increases in the level of taurine (1.94-fold), lysine (1.82-fold) and ornithine (1.71-fold) in 4-week-old JR5558 retinas, which are metabolites associated with cellular antioxidant stress and tissue repair. By 8 weeks of age, metabolites associated with energy metabolism, including methionine (2.32-fold), α-ketoglutarate (α-KG) (2.11-fold), malate (2.02-fold), glycerate (1.57-fold) and hypoxanthine (1.38-fold), were significantly increased in the JR5558 retinas compared to age-matched wild-type controls (Figure 3F-G). These results highlight the dynamic metabolic reprogramming associated with this model.

**Figure 3:**
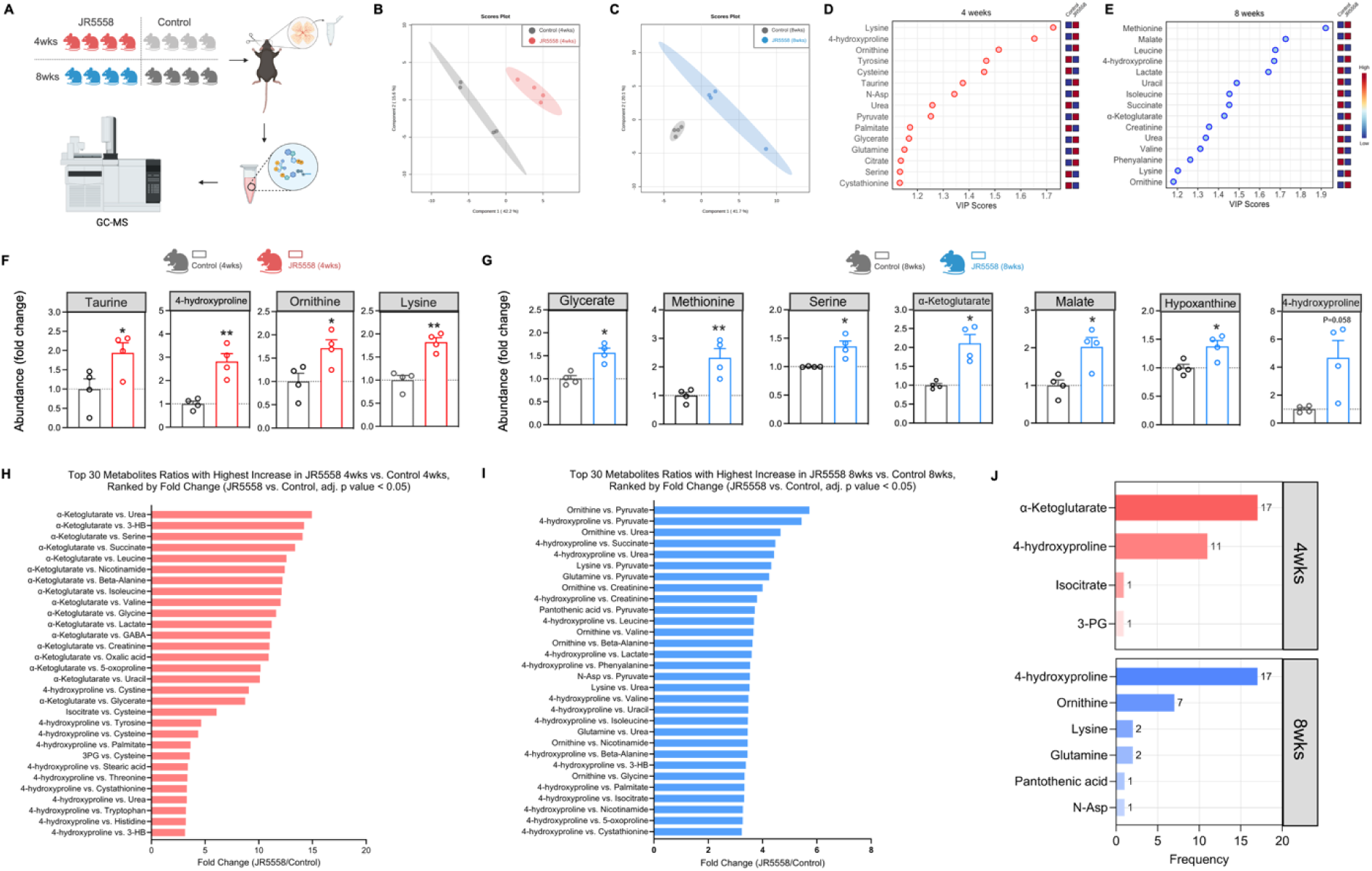
4-hydroxyproline was increased in the retinas of JR5558 mice. **(A)** Schematic diagram of experimental design of GC-MS based quantitative metabolomics in JR5558 and age-matched control retinas. **(B-C)** Partial Least Squares Discriminant Analysis (PLS-DA) analysis of JR5558 and control retinas at 4 weeks **(B)** and 8 weeks of age **(C). (D-E)** Top 15 metabolites with the highest VIP scores from 4-**(D)** and 8-week-old **(E)** JR5558 and control retinas. **(F-G)** Significantly increased metabolites at JR5558 retinas at 4 weeks **(F)** and 8 weeks **(G)** of age compared to control retinas. Data were shown as relative ion abundance over the control retinas. wks: weeks. Statistical analysis was performed using independent t test. n = 4. *: p < 0.05, **: p < 0.01, ***: p < 0.001, ****: p < 0.0001. All data are presented as means ± SEM. **(H-I)** The top 30 significant ratios with the highest increase ranked by fold change in JR5558 versus control retinas at 4 weeks **(H)** and 8 weeks of age **(I)**. **(J)** Frequency of numerators among the top 30 significant ratios in JR5558 and control retinas at both ages.

### Activation of P4HA1 in subretinal fibrotic lesions

4-hydroxyproline is derived from post-translational hydroxylation of proline residues in collagen peptides, a process catalyzed by collagen P4H, an Fe (II)- and α-ketoglutarate (αKG)-dependent enzyme (FAKGDs) [51]. P4H functions as an α2β2 heterotetramer, consisting of one of three catalytic α-subunits (encoded by *P4HA1*, *P4HA2* and *P4HA3*) and a structural β-subunit, protein disulfide isomerase (PDI, encoded by *P4HB*). Each α-subunit can combine with the β-subunit to form three tetrameric complexes: P4H1, P4H2 and P4H3 (Figure 4A) [52]. Since P4H is the enzyme responsible for the post-translational hydroxylation of proline residues in collagen to 4-hydroxyproline, we examined the transcriptional expression of all P4H subunits in the RNA-seq data from JR5558 and wild-type retinas. Aligning with the enrichment of arginine and proline metabolism in overlapped genes (Figure 2K), we found that only *P4ha1* mRNA was significantly different, being significantly lower at 4 weeks (Figure 4B) and significantly higher at 8 weeks (Figure 4C) in the JR5558 retinas.

**Figure 4:**
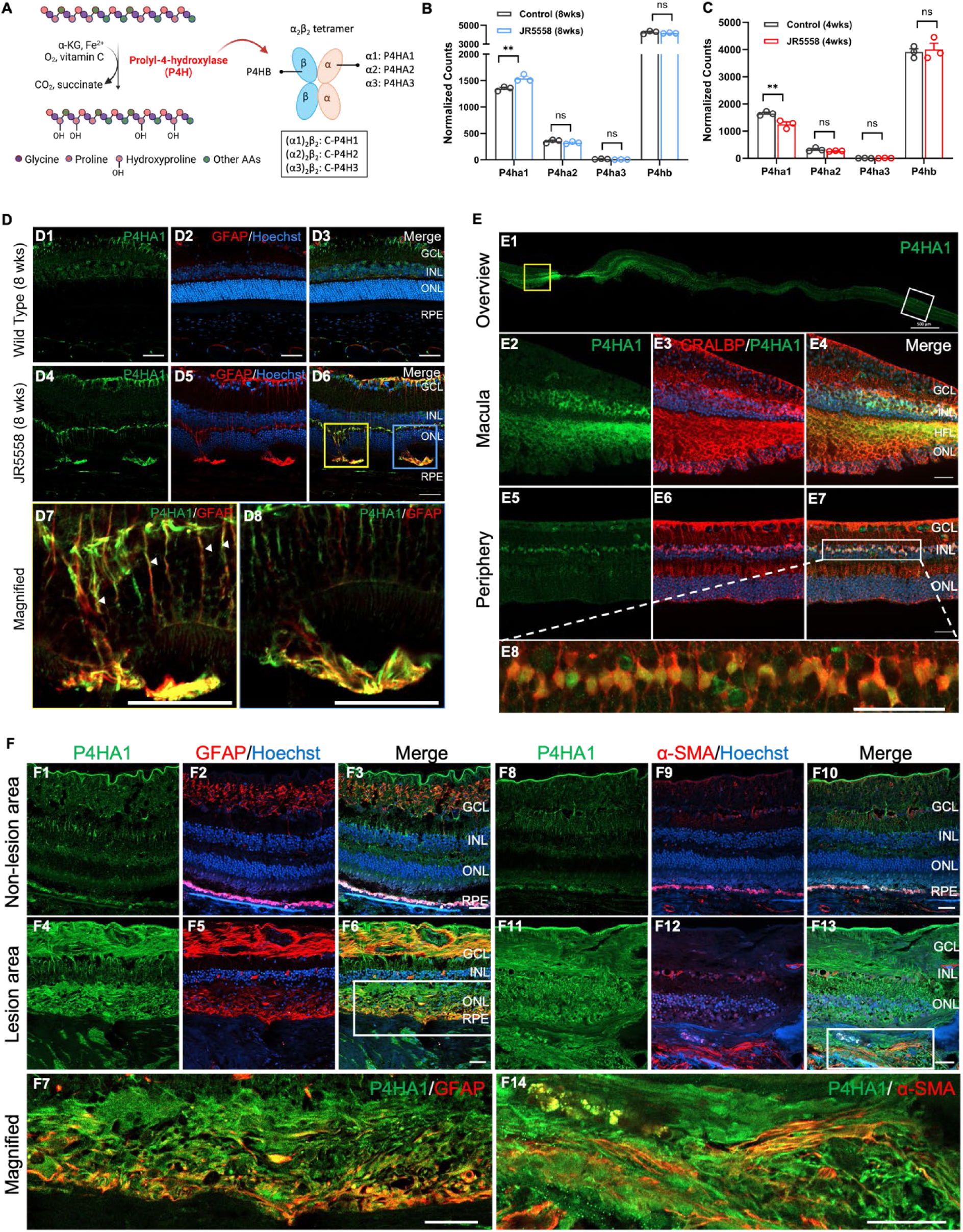
Increased expression of P4HA1 in the retinas of JR5558 and human AMD eyes. (A) Schematic of the enzymatic process of proline hydroxylation in collagen peptide (left). Structure of prolyl-4-hydroxylase as a heterotetramer was illustrated (right). **(B-C)** Normalized counts of P4ha1, P4ha2, P4ha3 and P4hb mRNA from JR5558 retinas were extracted from RNA-sequencing analysis. n= 3. **: p < 0.01, ns: not significant. All data are presented as means ± SEM. **(D)** Representative immunostaining of P4HA1 (green) and GFAP (red) in the retinas of 8-week-old control **(D1-D3)** and JR5558 **(D4-D8)** mice. Field-enlarged images from yellow box and blue box in **(D6)** were shown in **(D7)** and **(D8)**, respectively. Nuclei were stained with Hoechst (blue). Arrows in **(D7)** indicated discrete P4HA1 activation along Müller cell processes. Scale bar: 50 µm. **(E)** Representative immunostaining of P4HA1 (green) and CRALBP (red) in normal human retina on vibratome sections of the macula **(E2-4)** and periphery **(E5-8)**. Field-enlarged image from **(E7)** was shown in **(E8)**. Nuclei were stained with Hoechst (blue). Scale bar: 50 µm. **(F)** Representative immunostaining of P4HA1 (green) and GFAP/α-SMA (red) in nAMD patients with subretinal fibrosis on frozen sections of the non-lesion **(F1-3, F8-10)** and lesion area **(F4-6, F11-13)**. Field-enlarged image from the white box in **(F6)** and **(F13)** were shown in **(F7)** and **(F14)**, respectively. Nuclei were stained with Hoechst (blue). Scale bar: 50 µm. 8 wks: 8 weeks of age. GCL: ganglion cell layer. INL: inner nuclear layer. HFL: Henle’s fiber layer. ONL: outer nuclear layer. RPE: retinal pigment epithelium.

We first explored the localization of the expression of P4HA1 in 8-week-old JR5558 mice and age-matched control mice by immunofluorescent staining with P4HA1 and glial fibrillary acidic protein (GFAP), a marker for activated Müller cells and subretinal fibrosis[53]. Compared to control mice, strong activation of P4HA1 was observed in the GFAP-positive subretinal lesions of JR5558 mice. Notably, discrete P4HA1 activation was also observed in the RPE layer and along Müller cell processes (Figure 4D). Similar activation of P4HA1 was also seen around fibronectin-positive areas in mice eyes receiving the two-stage laser treatment, another model of subretinal fibrosis (**Figure S3**), confirming the involvement of P4HA1 in subretinal fibrotic microenvironment.

The staining of P4HA1 in vibratome sections of a normal human macula and peripheral retina is shown in Figure 4E. An overview of a long retinal slice showed stronger expression in the macula than in the peripheral retina, particularly in the Henle’s fiber layer (**Figure 4E1**). P4HA1 co-localized with CRALBP in both regions, indicating it was predominantly expressed in Müller cell bodies and processes in the retina (**Figure 4E2-8**). We then studied expression of P4HA1 and markers of fibrosis, GFAP and α-SMA, in cryosections of human eyes with nAMD-related subretinal fibrosis. Representative images from macular regions with or without fibrotic scars were shown in Figure 4F. P4HA1 was primarily expressed in RPE and inner nuclear layer of unaffected area. Extensive P4HA1 activation was observed in the GFAP-positive subretinal area, the ganglion cell layer and Müller cell processes (**Figure 4F1-7**). Expression of P4HA1 also co-localized with expression of α-SMA in the subretinal lesions (**Figure 4F8-14**).

### Inhibition of P4HA1 attenuated subretinal fibrosis in JR5558 mice

To explore the therapeutic potential of targeting P4HA1 for the treatment of subretinal fibrosis, we used a small molecule, diethyl pythiDC, to inhibit P4HA1 [32]. The AlamarBlue assay was performed to assess the safety of diethyl pythiDC in huRPEs and huPMCs after 24 hours. Diethyl pythiDC significantly inhibited cellular metabolic activity at 100 μM in huPMCs (Figure 5A), while no significant inhibition was found in huRPEs (Figure 5B). It has been reported that 10 μM is the highest safe dosage for clinical use and that diethyl pythiDC can inhibit P4H activity by 70% at this dose [32]. 10 μM diethyl pythiDC was thus chosen for further studies.

**Figure 5:**
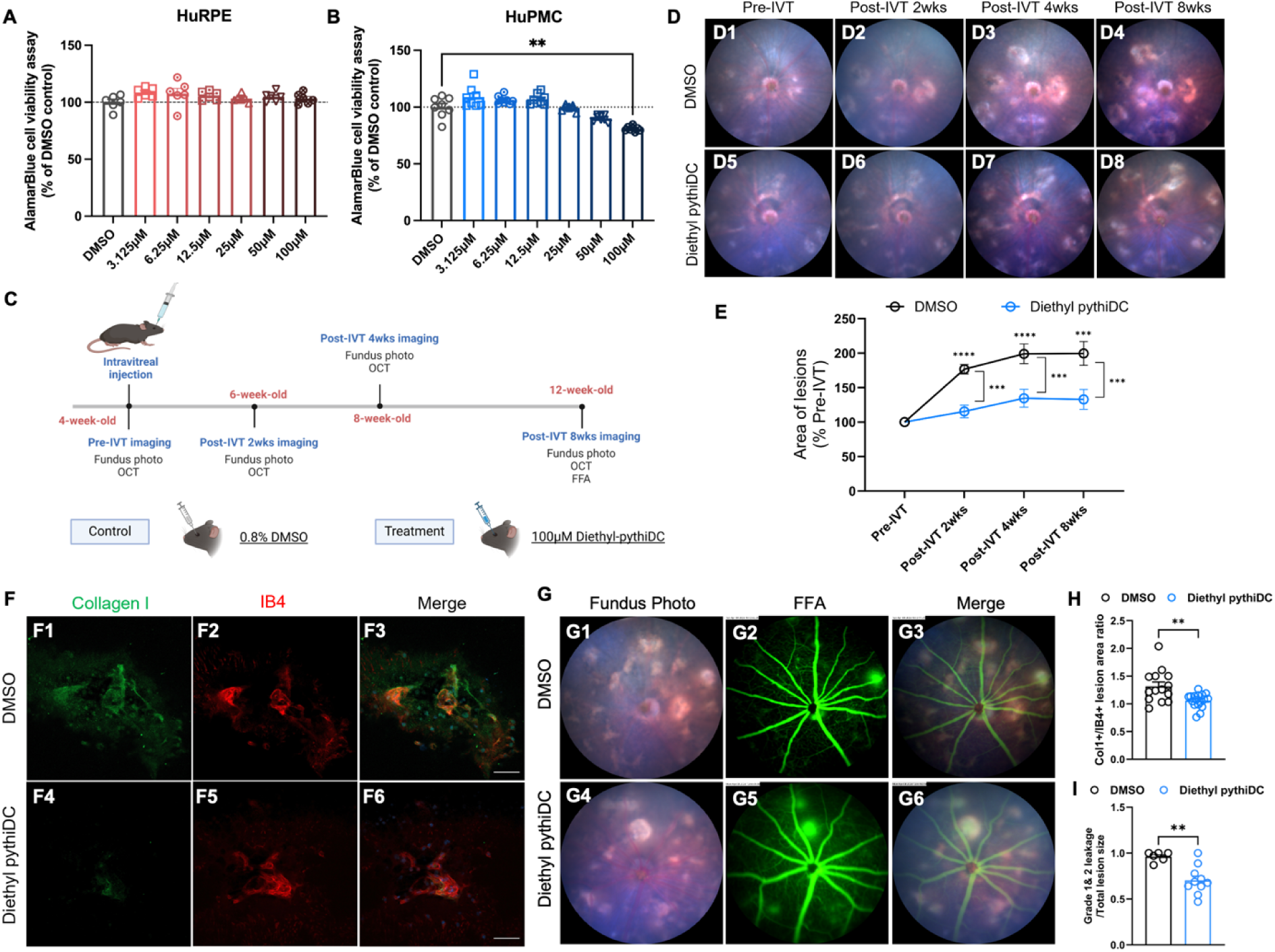
A P4H inhibitor, diethyl pythiDC, efficiently inhibited the development of fibrotic lesions in JR5558 mice. **(A-B)** AlamarBlue assay of huRPEs **(A)** and huPMCs **(B)** treated with different concentrations of diethyl pythiDC for 24 hours. Statistical analysis was performed using one-way ANOVA followed by Dunnett’s multiple comparison test. n = 6. **: p < 0.01. All data are presented as means ± SEM. **(C)** Experimental design schematic of intravitreal injection of JR5558 mice with 0.8% DMSO and 100µM Diethyl pythiDC (both were shown at working concentrations). **(D)** Representative images of fundus lesion areas in the DMSO control **(D1-4)** and diethyl pythiDC groups **(D5-D8)** at different timepoints. **(E)** Quantitative results of fibrotic lesions in DMSO-treated and diethyl pythiDC-treated groups. Data are normalized to pre-injection (pre-IVT) values. *Indicates a statistically significant difference compared to pre-injection baseline or DMSO control group. Statistical analysis was performed using two-way repeated measures ANOVA followed by Bonferroni multiple comparison test. n = 14-15. *: p < 0.05, **: p < 0.01, ***: p < 0.001, ****: p < 0.0001. **(F)** Representative immunostaining with IB4 and Collagen I in RPE/choroidal flatmounts from DMSO **(F1-3)** and diethyl pythiDC **(F4-6)** treated JR5558 mice 8 weeks post-IVT. Quantitative results of Col1+/IB4+ area ratios were shown in **(H)**. Statistical analysis was performed using independent t test. n = 14-17. **: p < 0.01. **(G)** Representative images of fundus photos and FFA from DMSO **(G1-3)** and diethyl pythiDC **(G4-6)** treated JR5558 mice to visualize primarily “fibrotic lesions” at 8 weeks post-IVT. Quantitative results were shown in **(I)**. Statistical analysis was performed using independent t test. n = 6-10. **: p < 0.01. All data are presented as means ± SEM.

We assessed the effect of intravitreal injection of 1 µL of 100 µM either diethyl pythiDC or 0.8% DMSO vehicle control (estimated final concentration in the vitreous: 10 µM or 0 µM diethyl pythiDC in 0.08% DMSO) into the eyes of 4-week-old JR5558 mice. Fundus imaging and OCT were performed before injection and at 2-, 4- and 8-weeks after the injection. FFA was included at 8 weeks post-injection (post-IVT) (Figure 5C). We found significantly fewer yellow-white lesions in fundus photos at 2-, 4- and 8-weeks post-IVT in diethyl pythiDC-treated group than in DMSO-treated controls (Figure 5D-E). RPE/choroid flatmounts from 8 weeks post-IVT were stained with IB4 and collagen I. As has previously been used to evaluate the degree of angio-fibrotic switch in preclinical studies, Col1+/IB4+ area ratio from each lesion was calculated [54]. Diethyl pythiDC-treated eyes showed a significantly lower mean Col1+/IB4+ ratio than DMSO-treated controls (Figure 5F**, H**), suggesting that diethyl pythiDC inhibited the development of fibrosis. Analysis of fundus photography and FFA also suggested that diethyl pythiDC inhibited the development of primarily fibrotic lesions 8 weeks after the injection (Figure 5G**, I**).

### Combination of diethyl pythiDC with aflibercept inhibited subretinal fibrosis more than monotherapies in Cluster 2 JR5558 mice

Many patients receiving VEGF inhibitors for subretinal neovascularization eventually lose vision from subretinal fibrosis[4, 5, 55]. Fibrotic lesions in nAMD patients originate from VEGF-induced vascular leakage and angiogenesis, but they develop even in patients treated with VEGF inhibitors [56, 57]. This suggests that adding anti-fibrotic drugs to VEGF inhibitors may provide a greater therapeutic benefit in some cases. To test this hypothesis, we intravitreally injected 1µL of 100µM diethyl pythiDC, 2.5µg/µL aflibercept, 2.5µg/µL aflibercept combined with 100µM Diethyl pythiDC, or 0.8% DMSO in 4-week-old JR5558 mice (Figure 6A). Fundus imaging was performed at 2, 4, and 8 weeks after the injection. Compared to DMSO-treated controls, diethyl pythiDC alone significantly reduced the lesion sizes at 2-, 4- and 8-weeks post-IVT, whereas aflibercept alone or in combination with diethyl pythiDC significantly reduced lesion size only at 2 weeks post-IVT (Figure 6B).

**Figure 6:**
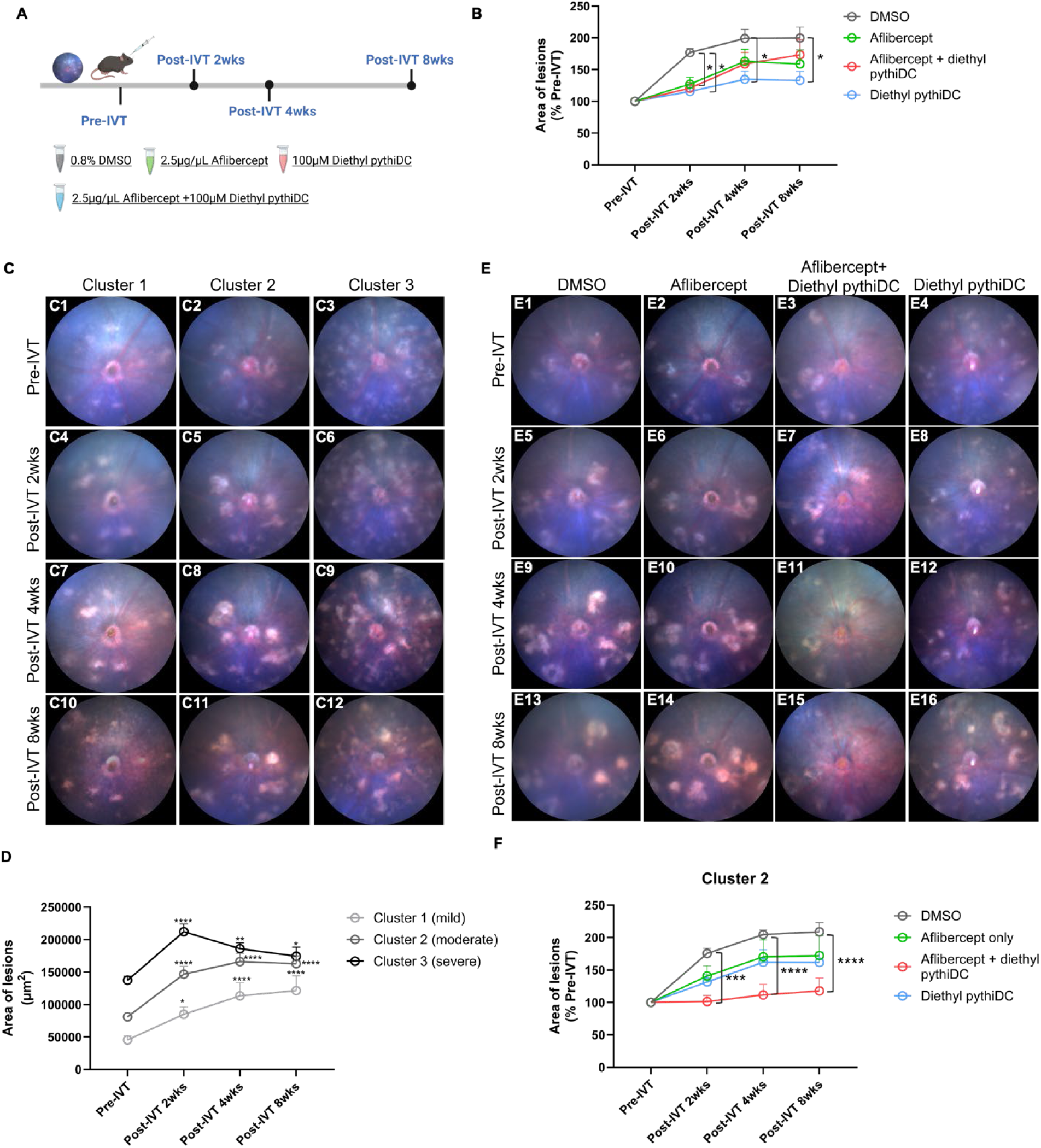
Combination of diethyl pythiDC with aflibercept efficiently inhibited the development of subretinal fibrosis in Cluster 2 JR5558 mice. **(A)** Experimental design schematic of intravitreal injection of JR5558 mice with 0.8% DMSO, 2.5µg/µL aflibercept, 2.5µg/µL aflibercept combined with 100µM Diethyl pythiDC, or 100µM Diethyl pythiDC (all were shown at working concentrations). **(B)** Quantification results of fundus lesion areas in all mice at different timepoints. *Indicates a statistically significant difference compared to DMSO control group at each timepoint. wks: weeks. Statistical analysis was performed using two-way repeated measures ANOVA followed by Tukey’s multiple comparison test. n = 14-19. **(C-D)** Representative images **(C)** and quantification results **(D)** of fundus lesion areas in 0.8% DMSO group of three clusters at different timepoints. *Indicates a statistically significant difference compared to pre-injection baseline. wks: weeks. Statistical analysis was performed using two-way repeated measures ANOVA followed by the Tukey’s multiple comparison test. n = 4-7. **(E-F)**Representative images **(E)** and quantitative results **(F)** of fundus lesion areas in Cluster 2 mice injected with 0.8% DMSO, 2.5µg/µL aflibercept, 2.5µg/µL aflibercept combined with 100µM diethyl pythiDC, 100µM diethyl pythiDC at different timepoints. *Indicates a statistically significant difference compared to DMSO control group at each timepoint. Statistical analysis was performed using two-way repeated measures ANOVA followed by Tukey’s multiple comparison test. n = 6-9. Pre-IVT: pre-injection. Post-IVT 2wks: 2 weeks post-injection. Post-IVT 4wks: 4 weeks post-injection. Post-IVT 8wks: 8 weeks post-injection. *: p < 0.05, **: p < 0.01, ***: p < 0.001, ****: p < 0.0001. Data are normalized to pre-IVT values. All data are presented as means ± SEM.

The sizes of subretinal fibrosis lesions at baseline in the DMSO-treated control group ranged from 28,344 μm² to 153,516 μm² (**Figure S4A**). To investigate whether the lesion sizes at baseline potentially affect the lesion progression, a K-means algorithm was applied to classify the baseline lesions in the DMSO-treated group into three clusters (k=3): 28,344–59,696 μm² for Cluster 1 (mild), 67,248–104,176 μm² for Cluster 2 (moderate) and 125,265–153,516 μm² for Cluster 3 (severe) (**Figure S4B**). Analysis of total lesion sizes up to 8 weeks after the injection of DMSO revealed distinct growth trajectories across three clusters. Subretinal lesions grew significantly at 2 weeks post-IVT in all clusters. While lesions in Cluster 1 and 2 continued to expand until 8 weeks post-IVT, those in Cluster 3 did not change in size significantly, reaching only 127% of their pre-IVT area at 8 weeks post-IVT (Figure 6C-D). Given the distinct growth trajectories across baseline lesion size clusters, we next assessed the efficacy of different treatments within each predefined cluster. In Cluster 2, characterized by moderate baseline lesion size and progressive lesion growth, the combination of aflibercept and diethyl pythiDC significantly suppressed lesion expansion at all post-IVT time points, outperforming monotherapies alone (Figure 6E-F).

### Diethyl pythiDC inhibited collagen production, secretion and contraction *in vitro*

We investigated the direct impact of diethyl pythiDC on collagen formation by assessing its ability to inhibit collagen turnover in huRPEs, a widely accepted cellular contributor to macular fibrosis *in vitro* [58]. Western blot analysis of huRPEs cell lysates revealed significant upregulation of collagen I and P4HA1 proteins, along with fibrotic markers fibronectin and α-SMA 3 days after treatment with 12.5 ng/mL TGFβ2 (Figure 7A-B). We then treated TGFβ2-stimulated huRPEs with 10 µM diethyl pythiDC or vehicle for 3 days to determine whether P4HA1 influences collagen turnover. Procollagen I is assembled and processed intracellularly before being secreted into the extracellular space, where it is cleaved into mature collagen fibrils along with C-propeptides and N-propeptides [59, 60]. Thus, proteins from both cell supernatants (Sup) and cell lysates (Lys) were extracted to detect procollagen I synthesis and secretion (Figure 7C). As shown in Figure 7D, TGFβ2 significantly increased procollagen I synthesized in cell lysates (Figure 7E), and procollagen I (Figure 7G) and C-propeptides (Figure 7H) secreted into the medium, all of which could be inhibited by diethyl pythiDC treatment. Of note, addition of diethyl pythiDC alone did not affect the protein level of P4HA1 compared to controls, whereas it inhibited the increase in P4HA1 that was induced by TGFβ2 (Figure 7F). TGFβ2-stimulated huRPEs were seeded in 3D collagen gels and treated with diethyl pythiDC or vehicle from day 0 to day 3. As shown in the representative images (Figure 7I), all groups exhibited the fastest contraction rate on day 1, with the gel size reduced to 57%, 39% and 48% of Day 0 in the control, TGFβ2-treated, TGFβ2 and diethyl pythiDC-treated groups, respectively. The contraction rate subsequently began to slow down after day 1 (Figure 7J). For intergroup comparison, the gel size of each group at each time point was normalized to the control group. The gel size of TGFβ2-treated cells was significantly smaller than that of vehicle controls, whereas the addition of Diethyl pythiDC reversed this contraction effect, with gel sizes comparable to those of the control group (Figure 7K).

**Figure 7:**
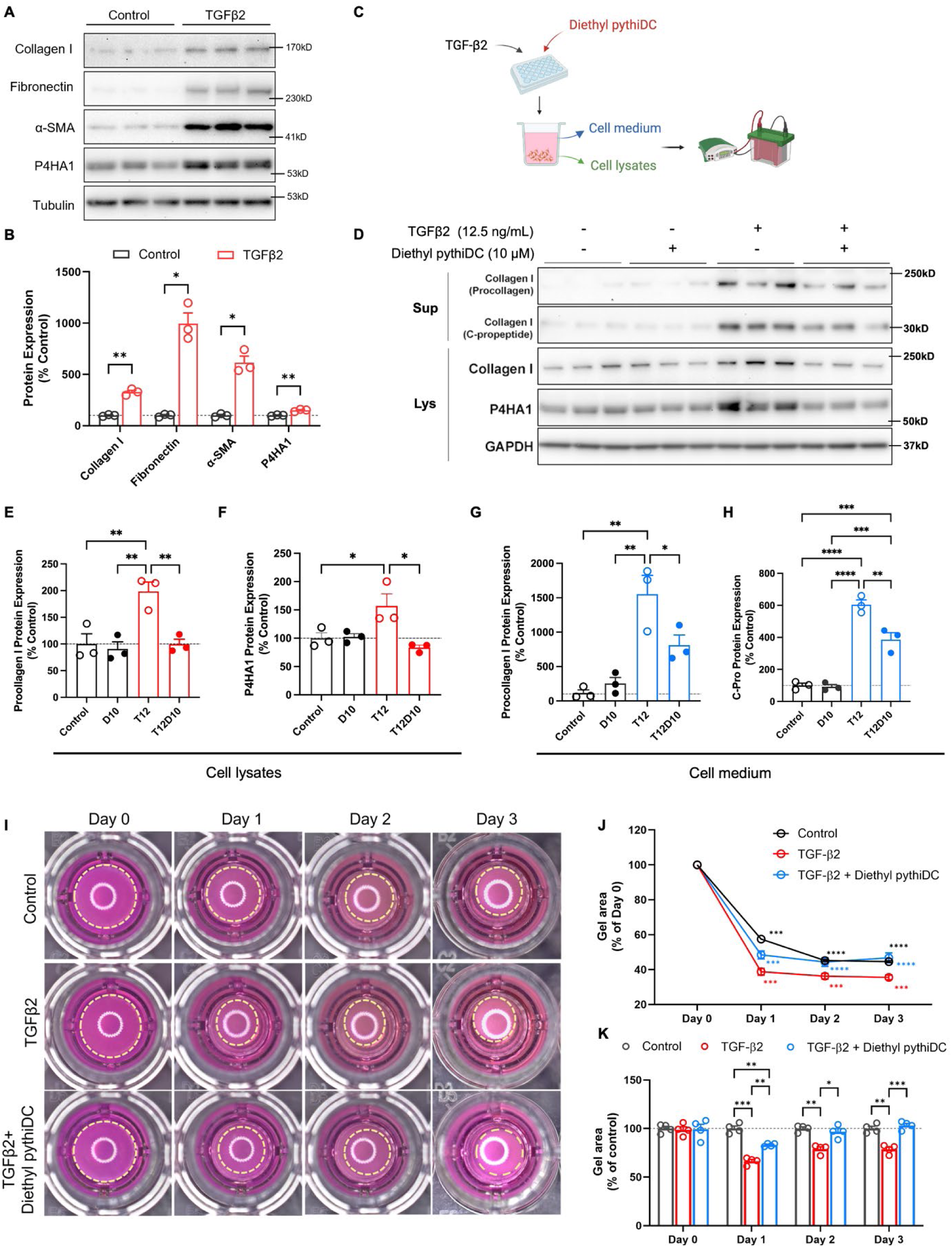
Diethyl pythiDC inhibited collagen synthesis, production and contraction in huRPEs. **(A-B)** Representative images **(A)** and quantification results **(B)** of Western blot in TGFβ2-treated and control huRPEs. Statistical analysis was performed using multiple unpaired t test. n = 3. *: p < 0.05, **: p < 0.01. **(C)** Workflow schematic of cell and medium protein collections in huRPEs after TGFβ2 stimulation with or without diethyl pythiDC. **(D-H)** Western blot of Collagen I with different molecular weights in the medium and cell lysates, along with P4HA1 in huRPEs after TGFβ2 stimulation with or without diethyl pythiDC for 3 days. Representative Western blot images from both medium (supernatant, Sup) and lysates (Lys) were shown in **(D)**. Quantification of procollagen I **(E)**, P4HA1 **(F)** in cell lysates and procollagen I **(G)**, C-propeptides **(H)** in cell medium were shown. T12: 12.5 ng/mL TGFβ2, D10: 10 μM diethyl pythiDC, T12D10: 12.5 ng/mL TGFβ2+10 μM diethyl pythiDC (all shown at final concentrations). Statistical analysis was performed using one-way ANOVA followed by Tukey’s multiple comparison test. n = 3. *: p < 0.05, **: p < 0.01, ****: p < 0.0001. **(I-K)** Representative images **(I)** of collagen gel contraction assay and quantitative results **(J-K)**. HuRPEs were treated with TGFβ2 and diethyl pythiDC for up to 3 days. Images for each collagen gel was taken every day. Gel areas illustrated by yellow dotted circles in **(I)** were calculated for each group. Statistical analysis was performed using two-way repeated measures ANOVA followed by Dunnett’s multiple comparison test in **(J)** and two-way repeated measures ANOVA followed by Tukey’s multiple comparison test in **(K)**. n = 3. *Indicates a statistically significant difference compared to Day 0 at each timepoint in **(J)**. *: p < 0.05, **: p < 0.01, ***: p < 0.001, ****: p < 0.0001. All data are presented as means ± SEM.

## Discussion

The biosynthesis of building blocks in collagen has emerged as a promising target for treating fibrotic diseases [61]. This study demonstrates the significant role of proline metabolism, particularly proline hydroxylation mediated by P4H, in the pathogenesis of subretinal fibrosis. Here, we have characterized the progressive development of fibrovascular lesions in JR5558 mice beginning as early as 4 weeks and significantly increasing by 8 weeks of age with integrated imaging, transcriptomic and metabolomic analyses. We found elevated levels of 4-hydroxyproline, along with significant changes in collagen-enriched extracellular matrix components, highlighting active collagen remodeling, which is a hallmark of fibrotic activity in this model. We also found the P4H subunit, P4HA1, was highly upregulated in fibrotic lesions in the two mouse models of subretinal fibrosis as well as in the eyes of patients with nAMD. The P4H inhibitor, diethyl pythiDC, effectively suppressed the development and progression of fibrosis in JR5558 mice. Combining diethyl pythiDC with aflibercept resulted in enhanced anti-fibrotic effects for the moderate phenotype, suggesting a synergistic therapeutic strategy. These findings highlight P4HA1 as a potential therapeutic target and underscore the potential for multi-targeted approaches to manage nAMD-associated fibrosis.

### Phenotypic and functional changes in the progression of subretinal fibrosis

The limited availability of animal models has restricted drug development and clinical management of subretinal fibrosis. Recent studies that focused on this issue have identified new animal models, including JR5558 transgenic mice [11, 53, 62–64]. By further studying the progression of fibrotic components in JR5558 mice, findings from our imaging analysis are consistent with a recent study that reported a significant increase of collagen remodeling, as measured by collagen hybridizing peptides (CHP), in 56-day-old JR5558 mice (Figure 1) [65].

Our transcriptomic study found that differential gene patterns associated with pro-fibrotic pathways occurred as early as 4 weeks of age in JR5558 mice, providing a molecular cue for determining the optimal treatment window in this model (Figure 2). Significant changes in inflammatory pathways, including the complement system, acute phase response and necroptosis signaling pathways, in JR5558 mice at 4 and 8 weeks of age suggest that altered immune responses are key features of subretinal fibrosis (Figure 2D-E). Similar findings have also been reported in several other animal models of subretinal fibrosis [50, 66–68]. Our recent study integrating machine learning, molecular imaging and transcriptomic analysis in JR5558 mice has supported this idea. This new approach has identified several key genes involved in the inflammatory process, such as Complement component 1, Phospholipase C and Chemokine ligand 12, offering potential therapeutic targets to reduce subretinal fibrosis [69]. Clinical studies have also found that nAMD patients with subretinal fibrosis had aberrant immune profiles, such as altered serum levels of complement fragments, higher inflammatory cytokines and lower CD4+ T cells [70–72]. Alterations in photoreceptor pathways were also evident in JR5558 mice, perhaps due to the disruption of outer retinal architecture by the robust inflammation. Pathological wound healing can thus be triggered at this stage by local inflammation and chronic injury, leading to excessive collagen-enriched ECM deposition in JR5558 retinas [73]. This idea has been supported by immunostaining showing disruption of cones and rods, as well as previous findings of increased microglia activation in the outer retina of JR5558 mice [63].

We have found that the JR5558 transgenic mouse is a useful model for subretinal fibrosis that can be used to study the early molecular drivers of this irreversible process through the interplay of multiple pathways. While it effectively replicates key aspects of subretinal fibrosis, it may not, however fully capture the human condition. A potential limitation is applying this model to age-related human disease, as both phenotypic and transcriptomic changes start to occur in young (4- and 8-week-old) mice. Despite this, this model closely mimics human nAMD through the spontaneous and progressive development of fibrovascular lesions. Although transcriptomic analysis in neural retinas of JR5558 retinas revealed various functional changes, an integrated RPE/choroid analysis would further enhance our understanding of the cellular interactions driving the progression of fibrosis.

### Collagen Metabolism and Remodeling in Subretinal Fibrosis

Collagen is a triple-helical structure composed of repeating Gly-X-Y sequences, where Gly represents glycine at every third amino acid residue. Proline and hydroxyproline account for 9-13% of all amino acids at X and Y positions [74]. Collagen remodeling, a hallmark of fibrotic diseases, has been previously reported in JR5558 RPE/choroidal flatmounts using CHP staining [65]. Elevated levels of 4-hydroxyproline, a known marker of active collagen turnover and, thus, a common readout for fibrotic activity, were detected in JR5558 retinas (Figure 3). This aligns with the significant enrichment of arginine and proline metabolic pathways in genes shared across both age groups in JR5558 mice (Figure 2J-K). These findings align with clinical observations that report increased hydroxyproline in the vitreous of patients with proliferative diabetic retinopathy (PDR) and rhegmatogenous retinal detachment, both characterized by extensive disruption of collagen [75].

Metabolic reprogramming was evident through alterations in other amino acids critical for collagen biosynthesis, in addition to 4-hydroxyproline, such as lysine and serine, in JR5558 retinas (Figure 3F-G) [59]. As blocking *de novo* serine/glycine biosynthesis has shown therapeutic potential in pulmonary fibrosis, it would be of interest to explore whether key enzymes in this process may be potential drug targets in subretinal fibrosis [76, 77]. Ornithine, a precursor for proline and key intermediate in the urea cycle, was also elevated in 4-week-old JR5558 retinas (Figure 3F). Preclinical studies have reported that the urea cycle (ornithine and arginine) contributes to collagen biosynthesis and thus pulmonary fibrosis, suggesting a possible role of ornithine metabolism in nAMD-related fibrosis [78, 79].

Dysregulation of collagen metabolism may contribute to active collagen remodeling, leading to the formation of subretinal fibrosis. Given the protective role of *de novo* serine biosynthesis for Müller cell survival, one should bear in mind that targeting this pathway could predispose the macula to oxidative stress and, therefore, retinal degeneration [41, 80]. The dynamic metabolic networks of lysine and serine also complicate their direct roles in collagen biosynthesis [81, 82]. Apart from those building blocks for collagen synthesis, the exact role of ornithine in collagen deposition and its potential crosstalk with proline hydroxylation in nAMD-related fibrosis still warrants further research.

### Involvement of P4HA1 in the development of subretinal fibrosis

Although clinical studies have linked P4H subunits with high myopia and a congenital-onset disorder of connective tissue [83–86], we believe that this is the first report of P4HA1 expression and its role in the retina. Based on KEGG analysis of overlapped genes, we identified a consistent role of proline metabolism and its key enzyme P4HA1 in JR5558 retinas at both ages (Figure 2J-K). The significant activation of P4HA1 in 8-week-old JR5558 retinas and two-stage laser mice retinas, particularly in subretinal lesions, reinforces the idea that enhanced prolyl hydroxylation by P4HA1 contributes to subretinal fibrosis (Figure 4D**, S3**). This pattern of activation was mainly found in Müller cells and RPE cells, in line with the concepts that these cells produce a wide range of ECM proteins to support the structure of the retina and contribute to fibrotic scars in response to retinal insult [58, 87, 88]. Remarkably, P4HA1 was highly expressed in macular Müller cells, particularly in Henle’s fiber layer, indicating a more active role of Müller cells in ECM remodeling in the macula and, thus, higher susceptibility to fibrotic reaction to neovascularization (Figure 4E). This idea was supported by P4HA1 activation in the fibrotic macular lesions in eyes from a patient with nAMD (Figure 4F).

*In vitro* studies have implicated TGFβ2 as a potential driver of P4HA1 protein upregulation in huRPEs (Figure 7A-B), expanding the findings of a recent study that reported upregulation of P4ha2, P4ha3, P4hab mRNAs in dedifferentiated RPE cells induced by pro-fibrotic cytokines and multiple passages[89]. Several mechanisms may contribute to P4HA1 activation driven by TGFβ2 in huRPEs. It could increase the binding of upstream stimulatory factors (USF) to the E-box sequence in the P4HA1 gene, enhancing its promoter activity as observed in HeLa cells[90]. Additionally, TGFβ2 may boost glutamine catabolism, providing more proline substrates and thereby enhancing P4HA1 enzymatic activity [91].

These findings highlight a potential role of P4HA1 in nAMD-related macular fibrosis. Notably, we observed an increase of 4-hydroxyproline accompanied by downregulation of P4HA1 in 4-week-old JR5558 mice (Figure 3F, **4B**). High mRNA or protein level are not necessarily a result of high enzyme activity. It has previously been reported that glycosylation at the N259 site enhanced P4HA1 enzyme activity, leading to a lower protein expression [92]. Whether a similar post-translational modification occures in 4-week-old JR5558 retinas still requires further investigation. While P4HA1 activation was observed in both Müller cells and RPE cells, the cell-specific role of P4HA1 in the progression of subretinal fibrosis remains unclear.

### Therapeutic potential of inhibiting P4H in subretinal fibrosis

Our data demonstrate that diethyl pythiDC directly suppresses collagen synthesis and deposition both *in vitro* (Figure 7) and *in vivo* (Figure 5) by disrupting the enzymatic activity of P4H. In the JR5558 mouse model, subretinal fibrosis is characterized by a cumulative buildup of collagen-rich lesions that peaksd between 4 and 8 weeks of age (Figure 12). Intravitreal injection of diethyl pythiDC during the early stage (at 4 weeks of age) effectively halted lesion progression, with rapid and sustained antifibrotic effects observed up to 8 weeks post-injection (Figure 5). These results suggest that inhibition of proline hydroxylation is sufficient to disrupt the fibrotic cascade at its foundational stage, prior to irreversible extracellular matrix remodeling.

We observed variably sized subretinal lesions in 4-week-old JR5558 mice (**Figure S4**), which has also been reported previously [37, 38, 46, 63]. Lesion size at baseline stratified the mice treated with DMSO into distinct patterns of lesion progression, with treatment efficacy varying accordingly (Figure 6C-D). This has important implications for preclinical drug screening using this model. Subretinal fibrosis in JR5558 mice progresses via an angio-fibrotic switch, marked by regression of neovascular components and expansion of fibrotic tissue, ultimately leading to a stable scar phenotype [93]. Lesions in “Cluster 2” — those with moderate baseline size — exhibited rapid and sustained enlargement without spontaneous regression, closely resembling the fibrotic progression seen in human nAMD [94]. The lesions in Cluster 1 remained relatively mild throughout follow-up so they may not provide a robust fibrosis phenotype suitable for therapeutic testing. The Graph Pseudotime Analysis based on transcriptomics data from the 23 JR5558 mice suggested that the treatment window in JR5558 mice appears narrow, as delayed intervention may miss the “point of no return”, after which critical molecular transition occurs and fibrosis may become treatment-resistant[95]. Cluster 3 lesions may have already exceeded the optimal therapeutic window and become less responsive to therapies than those in Cluster 1 and 2. The moderate disease state, Cluster 2, might offer an optimal phenotype for anti-fibrotic drugs screening. Notably, combined treatment with aflibercept and diethyl pythiDC yielded a durable inhibitory effect in Cluster 2 (Figure 6E-F), indicating the combined therapy as a promising therapeutic strategy to treat subretinal fibrosis in this moderate disease phenotype. Nevertheless, further studies with larger sample sizes within this lesion size range are warranted to validate the predictive value of this lesion stratification approach.

It is worth mentioning that diethyl pythiDC is a pan-P4H inhibitor, so we cannot be certain that P4HA1 is the direct target responsible for the promising results we observed. As no subunit-specific inhibitors are available, genetic manipulation would be required to confirm the underlying contributor definitively. The complex interactions among P4H isoforms make it harder to identify the specific target of diethyl pythiDC in our case. As previously reported, P4HA1 can largely compensate for the absence of P4HA2 [96]. P4HA1 knockdown in colorectal cancer cells was also shown to decrease the expression of P4HA2 and P4HB [33]. Thus, the interplay among P4H isoforms cannot be overlooked.

In addition to its direct role in collagen remodeling, P4HA1 has been reported to act as an immune, epigenetic or metabolic regulator, suggesting additional mechanisms by which P4HA1 may contribute to subretinal fibrosis. For example, P4HA1 could govern metabolic perturbation through cofactors α-KG and succinate, leading to HIF-1α stabilization and epigenetic remodeling[97–99]. Taking cues from their roles in the cancer field, other potential areas of action of P4HA1 include immune modulation, amino acid metabolism rewiring and the regulation of critical signaling pathways, such as Hippo-YAP pathway[20, 99, 100].

Our findings collectively indicate that proline hydroxylation is a significant contributor to the development of subretinal fibrosis. Further in-depth validation highlighted the therapeutic potential of P4HA1 inhibition using diethyl pythiDC as a novel treatment strategy. The synergistic effect of combining diethyl pythiDC with anti-VEGF therapy underscores the potential of multi-targeted approaches for managing nAMD-associated fibrosis. Stratifying JR5558 lesions by initial area could ensure a more reliable and translatable evaluation of drug efficacy in this model. Future research should prioritize the development of isoform-specific P4H inhibitors to optimize therapeutic outcomes. Mechanistic studies on P4HA1 activity are warranted, including whether P4HA1 upregulation is post-translationally modified, whether P4HA1 rewires amino acid metabolism, whether P4HA1 bridges immune responses to a fibrotic phenotype and how P4HA1 interacts with other P4H isoforms. Expanding investigations to other preclinical models and exploring patient-derived organoids will improve the translatability of these findings to phase 1 clinical trials.

## Acknowledgement

The authors thank New South Wales Tissue Bank and Australian Ocular Bank for providing human eye tissues. This study received financial support from National Health and Medical Research Council (Investigator Grant GNT_1195021 for MG, Ideas grant GNT_ 2020950 for LZ and EC), the Australian Vision Research (for MG, LZ and TZ), National Eye Institute (EY031324 and EY032462 for JD) and Retinal Research Foundation (JD).

## Author contributions

T.Z., J.D., L.Z., M.G., Y.Z. conceptualized and designed the study; Y.Z., T.Z., S.L., M.Y., J.D., Jialing Z., S.Z., M.E, X.W., J.Y., Meidong Z. carried out experiments; Y.Z., L.Z., T.Z., M.G., E.C., J.D., Jingwen Z, S.L. analyzed and interpreted the data; Y.Z., T.Z., L.Z. wrote the manuscript; M.G., E.C., J.D., A.C., H.C., Meixia Z., Meidong. Z. critically revised the manuscript. All authors approved the final version to be published.

