## Supplementary figures for "Dysregulated Proline Metabolism Contributes to Subretinal Fibrosis in Neovascular AMD: Therapeutic Potential of Prolyl-4-Hydroxylase Inhibition"

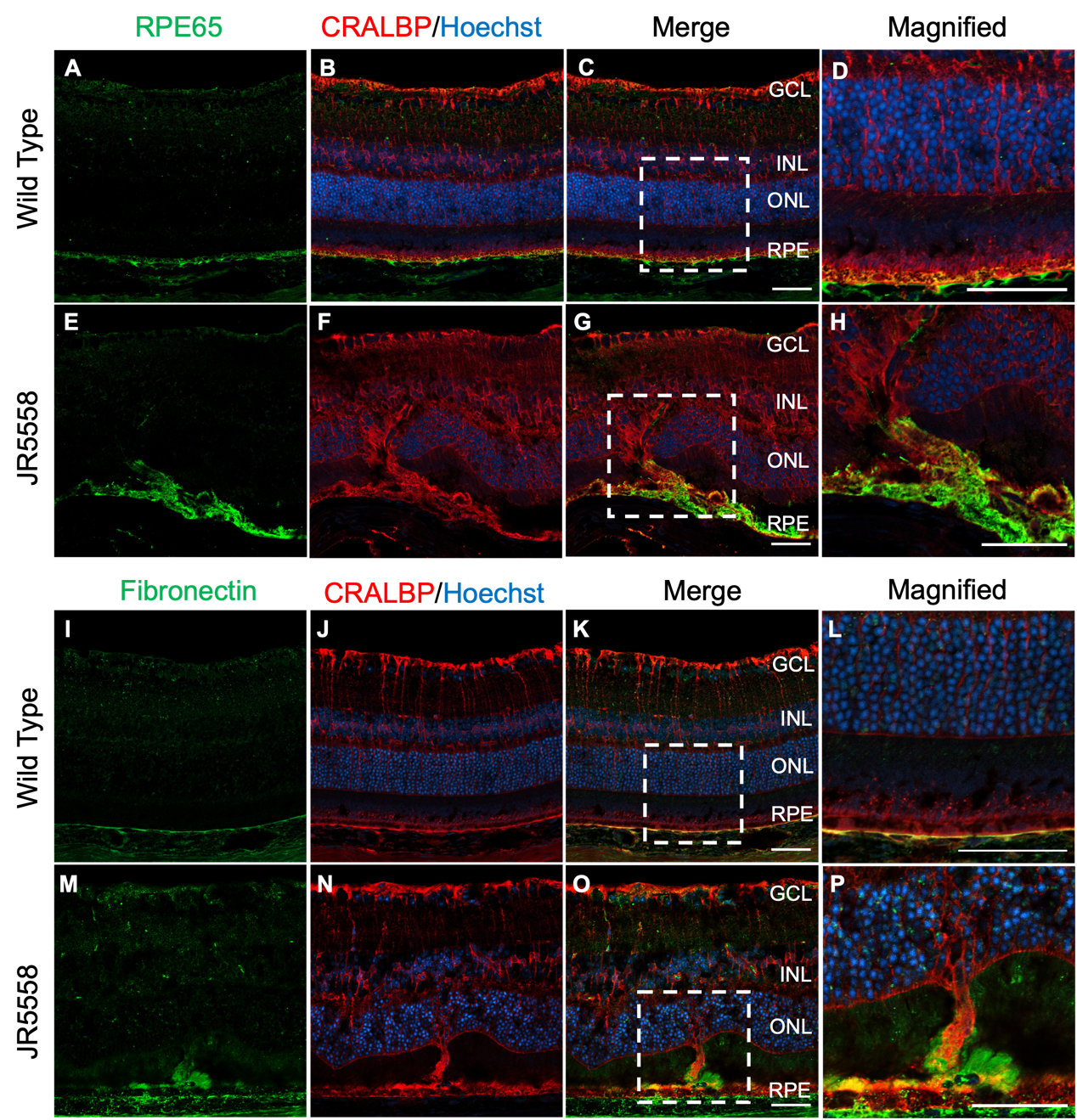

**Supplementary figure 1: Fibrovascular lesions in 8-week-old JR5558 mice could originate from Müller cells and RPE cells. (A-H)** Representative images of immunostaining of RPE65 (green) and CRALBP (red) in wild type mice **(A-D)** and JR5558 **(E-H)** retinas. Field-enlarged image from **(C)** and **(G)** were shown in **(D)** and **(H)**, respectively. **(I-P)** Representative images of immunostaining of Fibronectin (green) and CRALBP (red) in wild type mice **(I-L)** and JR5558 mice retinas (M-P). Field-enlarged images from white boxes in **(K)** and **(O)** were shown in **(L)** and **(P)**, respectively. GCL: ganglion cell layer. INL: inner nuclear layer. ONL: outer nuclear layer. RPE: retinal pigment epithelium. Nuclei were stained with Hoechst (blue). Scale bar: 50 µm.

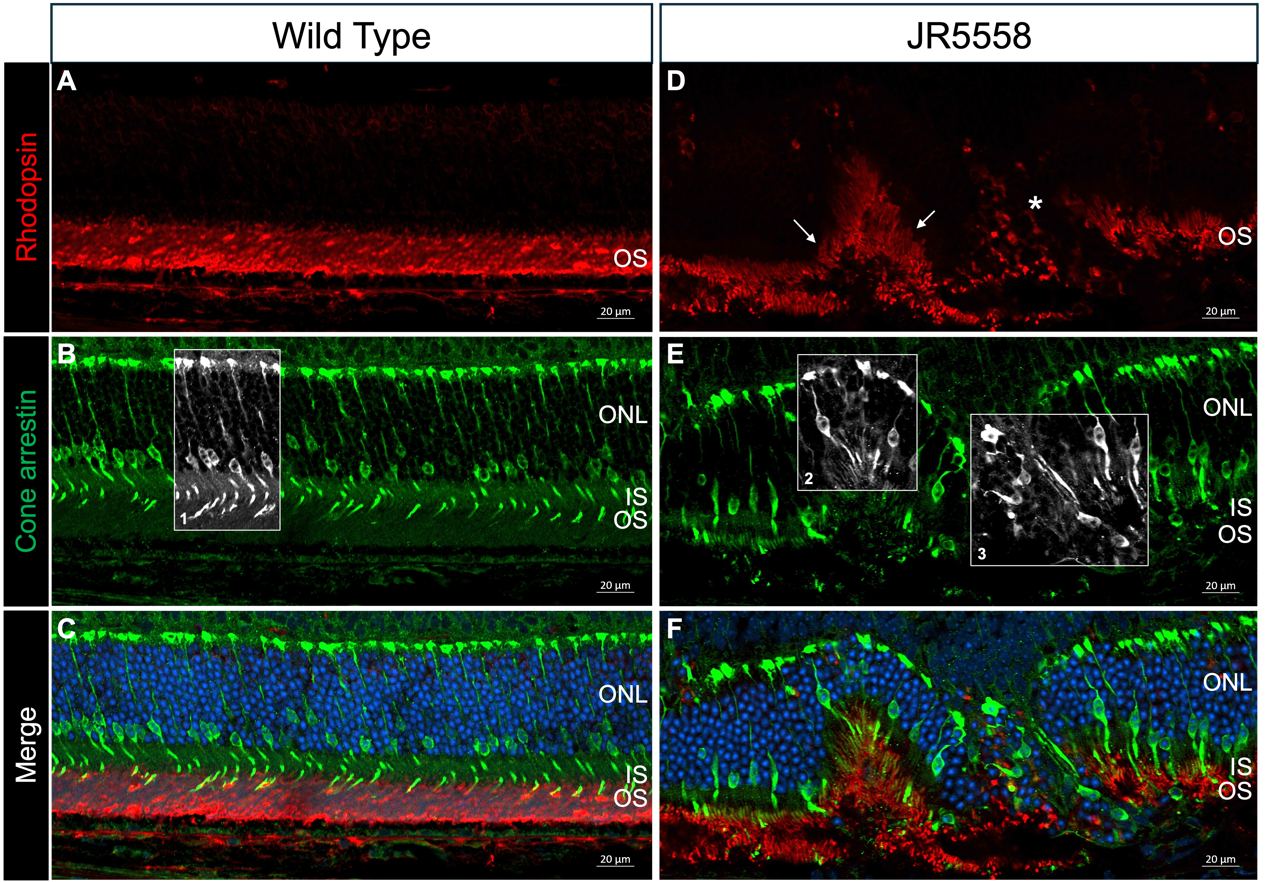

**Supplementary Figure 2: Photoreceptor damage in fibrovascular lesions in JR5558 retinas. (A-F)** Representative immunofluorescence images of retinal cryosections from wild-type mice **(A-C)** and JR5558 mice **(D-F)**. Region 1 in **(B)** showed an orderly arrangement of cone photoreceptors in wild type mice. Region 2 and 3 in **(E)** showed structural damage to cone photoreceptors in JR5558 mice. White arrow in **(D)** indicated disrupted rod photoreceptors, while the yellow asterisk highlighted the complete loss of rod outer segments. Red: rhodopsin. Green: cone arrestin. Nuclei were stained with Hoechst (blue). ONL: outer nuclear layer. IS: inner segment. OS: outer segment. Scale bar: 20 µm.

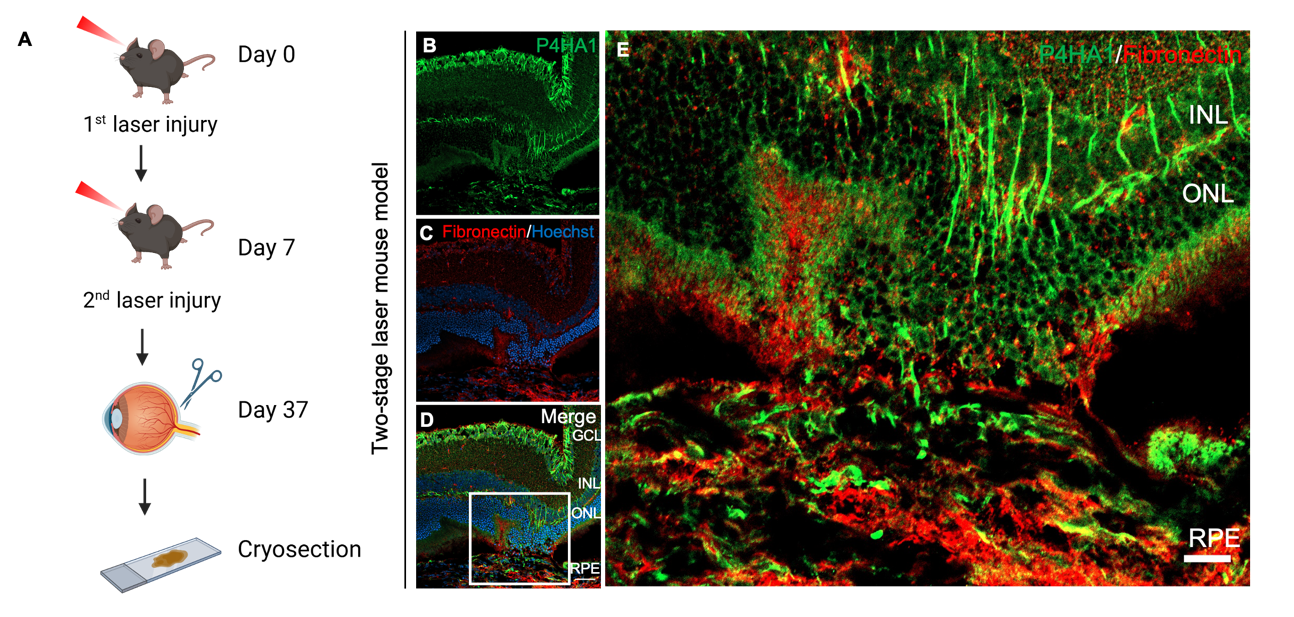

**Supplementary figure 3: Localization of P4HA1 in the subretinal lesion area in the two-stage laser mouse model. (A)** Experimental design schematic of the two-stage laser photocoagulation in mice. **(B-E)** Representative images of P4HA1 (green) and fibronectin (red) immunostaining on the cross-sections of retinas from C57BL/6 mice 30 days after the second laser photocoagulation. Field-enlarged image from the white box in **(D)** was shown in **(E)**. Nuclei were stained with Hoechst (blue). Scale bar: 50 µm. GCL: ganglion cell layer. INL: inner nuclear layer. ONL: outer nuclear layer. RPE: retinal pigment epithelium.

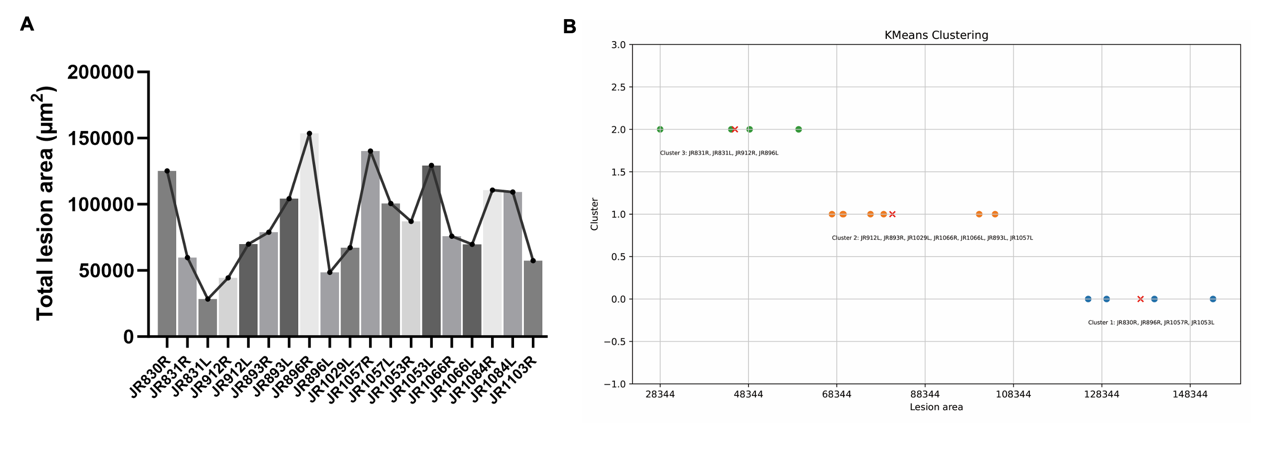

**Supplementary figure 4: Baseline distribution of fundus lesion sizes in JR5558 mice treated with 0.08% DMSO (final concentration).** Distribution of total lesion area **(A)** and K-means clustering results **(B)** of 0.08% DMSO treated JR5558 mice before the injection. Data in **(A)** are means ± SEM.
